# Large-genome phages encode diverse proteins associated with host cell envelope functions

**DOI:** 10.64898/2026.10.24.761521

**Authors:** Ling Yuan, LinXing Chen

## Abstract

Huge phages (or jumbo phages, >200 kilobases) encode extensive protein repertoires, yet their functional diversity and potential roles in host physiology remain poorly understood. Using protein structure-based analyses of dominant protein families across 7,187 non-redundant genomes of huge phages, we uncovered pronounced phylogenetic and habitat-associated diversification. Notably, we identified diverse proteins associated with bacterial cell envelope functions, particularly in human gut-associated huge phages. These included previously unreported phage-encoded sortase B, whose putative substrates contain signal peptides and cell-wall sorting signals, suggesting a potential route for displaying phage proteins on bacterial surfaces. Phage-encoded phage shock protein A (PspA) was distributed across diverse lineages and frequently neighbored genes associated with membrane organization and transport, implicating potential roles in membrane physiology. We also identified 1,603 β-barrel outer membrane proteins (OMPs) across 1,303 huge phages from diverse ecosystems. These OMPs exhibited structural diversity and associations with predicted host taxonomy, habitat, and neighboring envelope-related genes. Metatranscriptomic analysis supported expression of representative phage-encoded PspA and OMP genes in natural communities including the human gut. Together, these findings reveal bacterial cell envelope processes as a previously underappreciated potential interface for huge phage–host interactions.

## Introduction

Phages rely on their bacterial hosts to complete their infection cycles and have evolved diverse strategies to manipulate host physiology, for example, by redirecting DNA replication, transcription and translation, reprogramming central and nutrient metabolism, and counteracting host antiviral defense systems ^1–6^. The bacterial cell envelope represents a critical interface between phages and their hosts. It mediates phage adsorption and genome entry, while its controlled disruption is required for progeny release ^7–9^. Accordingly, phage proteins involved in receptor recognition ^10^, genome delivery ^11^, and host cell lysis ^9^ have been extensively characterized. Beyond these canonical infection processes, however, much less is known about the extent to which phages encode accessory proteins associated with the physiological functions of the host cell envelope ^12^.

Bacterial envelope systems mediate nutrient transport, membrane homeostasis, stress responses, and cell-surface interactions ^13–15^, raising the possibility that phages may exploit or modulate these processes during infection via encoding and expressing associated genes. Large-genome phages provide a particularly compelling system to explore such host-interaction functions. Tailed phages with genomes >200 kbp, commonly referred to as jumbo phages ^16^ or huge phages ^17^, have been detected across diverse ecosystems ^18–23^. Their expanded genomes encode unusually large protein repertoires and support complex infection strategies, including extensive replication and transcription machinery ^17^, sophisticated host takeover mechanisms ^24^, and in some lineages, nucleus-like compartments ^25–27^. These observations suggest that genome expansion provides phages with increased capacity to encode accessory functions that interact with host cellular physiology. Yet the functional repertoire enabled by this expanded coding capacity remains incompletely understood, particularly for processes associated with the bacterial cell envelope.

A major obstacle to resolving these functions is the extraordinary sequence diversity of phage proteins ^28,29^. Metagenomics has uncovered thousands of huge phages, but a substantial proportion of their proteins cannot be annotated using sequence-based searches ^30^. Protein structures are generally more conserved than sequences and can reveal functional relationships that are difficult to detect from sequence similarity alone. Recent advances in protein structure prediction and structural comparison therefore provide an opportunity to systematically investigate the poorly characterized functional repertoire of huge phages ^31–35^. Integrating structural annotation with phylogenetic distribution, genomic context, and environmental information can further distinguish broadly conserved viral functions from accessory functions potentially associated with particular hosts or ecological settings.

Here, we performed a structure-informed analysis of the dominant proteins encoded by 7,187 non-redundant huge phages collected across diverse ecosystems ^23^. We constructed a protein structure atlas to characterize their functional repertoire and found extensive phylogenetic and habitat-associated variation in accessory functions. We then focused on an unexpectedly diverse set of proteins associated with host cell envelope functions. These included sortase B and substrate modules, PspA embedded within diverse membrane-associated genomic modules, and a large repertoire of outer membrane β-barrel proteins associated with different host lineages and environmental settings. Metatranscriptomic analyses showed activity of representative membrane-associated genes in natural communities. Together, these findings expand the known functional repertoire of huge phages and suggest that the bacterial cell envelope represents an underappreciated interface for phage-encoded accessory functions during infection.

## Results

### HPSDB: the huge phage protein structure database

The Huge Phage Genome Collection (HPGC) is the most comprehensive repository of huge phage genomes to date, comprising genomes reported up to October 2024 ^23^. HPGC originally contained 7,230 non-redundant genomes, and 43 of them were further identified as non-viral ^36^. We determined the genetic code for the 7,187 remaining HPGC genomes (Methods), with 7,003 genomes using code 11, and 154 and 30 of them using code 15 and code 4, respectively (Supplementary Table 1). Prediction using the respective genetic code obtained a total of 2,500,924 protein-coding genes (Figure 1a). We performed CD-HIT clustering (≥90% identity) on them and obtained a total of 1,509,176 clusters. Of these, 162,625 clusters (∼10.8%) contained members detected in at least three genomes, and the representative proteins of these clusters were used for subsequent protein structure prediction. Approximately 70.5% (114,595 out of 162,625) of the predicted structures had an average pLDDT score of ≥70 (Figure 1b), collectively representing 671,099 proteins. Structural clustering using Foldseek grouped these protein structures into 52,425 structure clusters (Supplementary Table 2), of which 77.6% (40,701) were singletons. These 52,425 structure clusters constitute the basis of the Huge Phage Protein Structure DataBase (HPSDB), which represents the dominant protein structures of huge phages.

**Figure 1.**
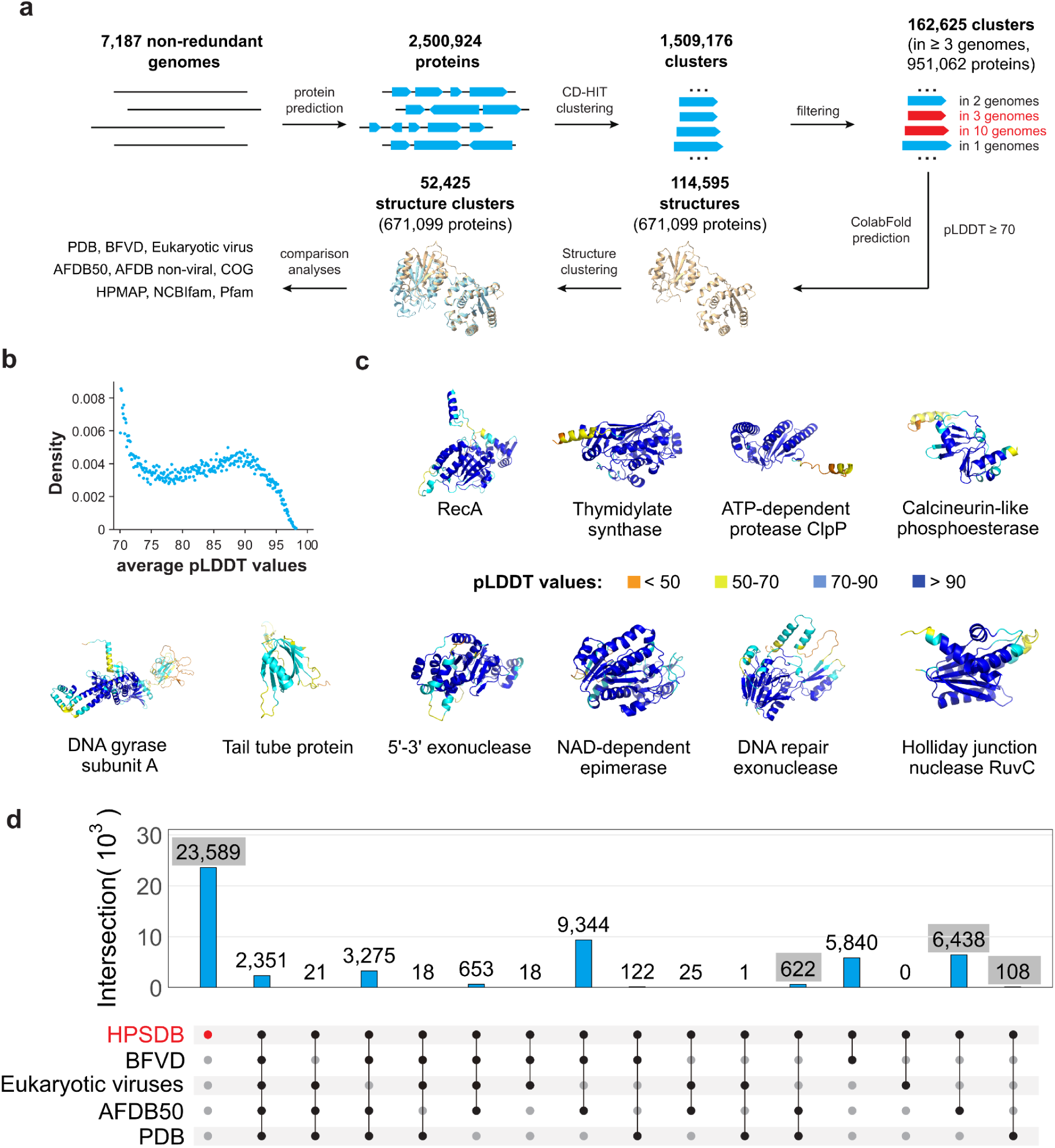
The huge phage protein structure database (HPSDB). (a) Workflow for HPSDB construction. The confirmed 7,187 non-redundant huge phage genomes in HPGC were used to predict protein-coding genes, which were then clustered using CD-HIT with a minimum identity of 90% (Methods). The representatives of CD-HIT protein clusters with members detected in at least 3 genomes were selected for protein structure prediction. The predicted structures with a minimum pLDDT value of 70 were clustered, and the representative structures were compared against those in protein structure references. (b) The distribution of the average pLDDT scores of the 114,595 HPSDB protein structures. (c) The functional annotation of the top 10 protein clusters with the most number of members in HPSDB. For each cluster representative, the pLDDT value ranges are indicated by different colors. (d) The shared and unique protein structures between HPSDB representatives and four reference structure databases, including BFVD (Big Fantastic Virus Database), the eukaryotic virus structure database, AFDB50, and PDB. The number of protein structures present in HPSDB, while absent in the other two virus-derived structure databases (i.e., BFVD and the eukaryotic virus structure database), is highlighted with a gray background.

To inform functional annotation, we searched HPSDB structure representatives against several reference protein structure databases, including BFVD (Big Fantastic Virus Database) ^37^, the eukaryotic virus structure database ^38^, AFDB50 ^39^, and Protein Data Bank (PDB) ^40^. As expected, the top 10 structure clusters with the most members were related to core functions of phages, including those related to DNA processing (RecA recombinase, DNA gyrase subunit A, 5’-3’ exonuclease, DNA repair exonuclease, Holliday junction nuclease RuvC), host manipulation mechanisms (Thymidylate synthase, ATP-dependent Clp protease, Calcineurin-like phosphoesterase, NAD-dependent epimerase), and virion assembly (Tail tube protein) (Figure 1c).

Notably, we identified 30,757 structural cluster representatives (accounting for 58.7% of all HPSDB structure clusters) that had no significant matches in the virus-derived reference database of BFVD or the eukaryotic virus structure database (Figure 1d). Among these, 23,589 also lacked significant matches in the databases AFDB50 or PDB, highlighting the substantial unexplored protein structural diversity encoded by huge phages.

### Broadly distributed and habitat-specific protein structure clusters in huge phages

Among the 7,187 huge phage genomes, the animal digestive system (primarily the human gut) represented the most prevalent environmental source, accounting for 2,216 genomes, followed by freshwater (1,881 genomes) and soil (929 genomes) (Supplementary Table 1). All 7,187 huge phage genomes were further clustered at ≥90% average nucleotide identity (ANI) and ≥85% alignment fraction (Methods), resulting in 4,223 genome clusters. The 90% clustering threshold was chosen to obtain a less redundant yet broadly representative set of genomes for downstream phylogenetic and protein-structure analyses rather than to define species-level vOTUs that typically require a 95% ANI threshold ^41^.

Based on HPSDB structural annotation and an additional round of iterative search (Methods), TerL (terminase large subunit) and MCP (major capsid protein) were identified in 3,095 (73.3%) and 3,019 (71.5%) out of the 4,223 genome clusters, respectively. Phylogenetic analyses of TerL (Figure 2a) and MCP (Supplementary Figure 1) revealed distinct clades associated with habitat origin and predicted bacterial host phyla. Mapping the presence–absence profiles of the abundant protein structure clusters (those detected in ≥ 20 genome clusters) to the TerL phylogeny showed distinct distribution patterns. Bacillota (previously Firmicutes) potentially infecting huge phages, which were primarily from the animal digestive system, formed independent phylogenetic clades and exhibited prominent modular protein organization (indicated by three dark red boxes, Figure 2a). In contrast, those predicted to target Bacteroidota and Pseudomonadota residing in clades with origins from multiple habitats, including the animal digestive system, shared protein structure sets across ecosystems (indicated by two brown boxes, Figure 2a). Huge phages from aquatic ecosystems (freshwater and marine) also showed multiple modular protein blocks aligned with specific host ranges (indicated by five blue boxes, Figure 2a). These findings suggested an ecological partitioning of huge phage functional repertoires that is associated with habitat origin and host range.

**Figure 2.**
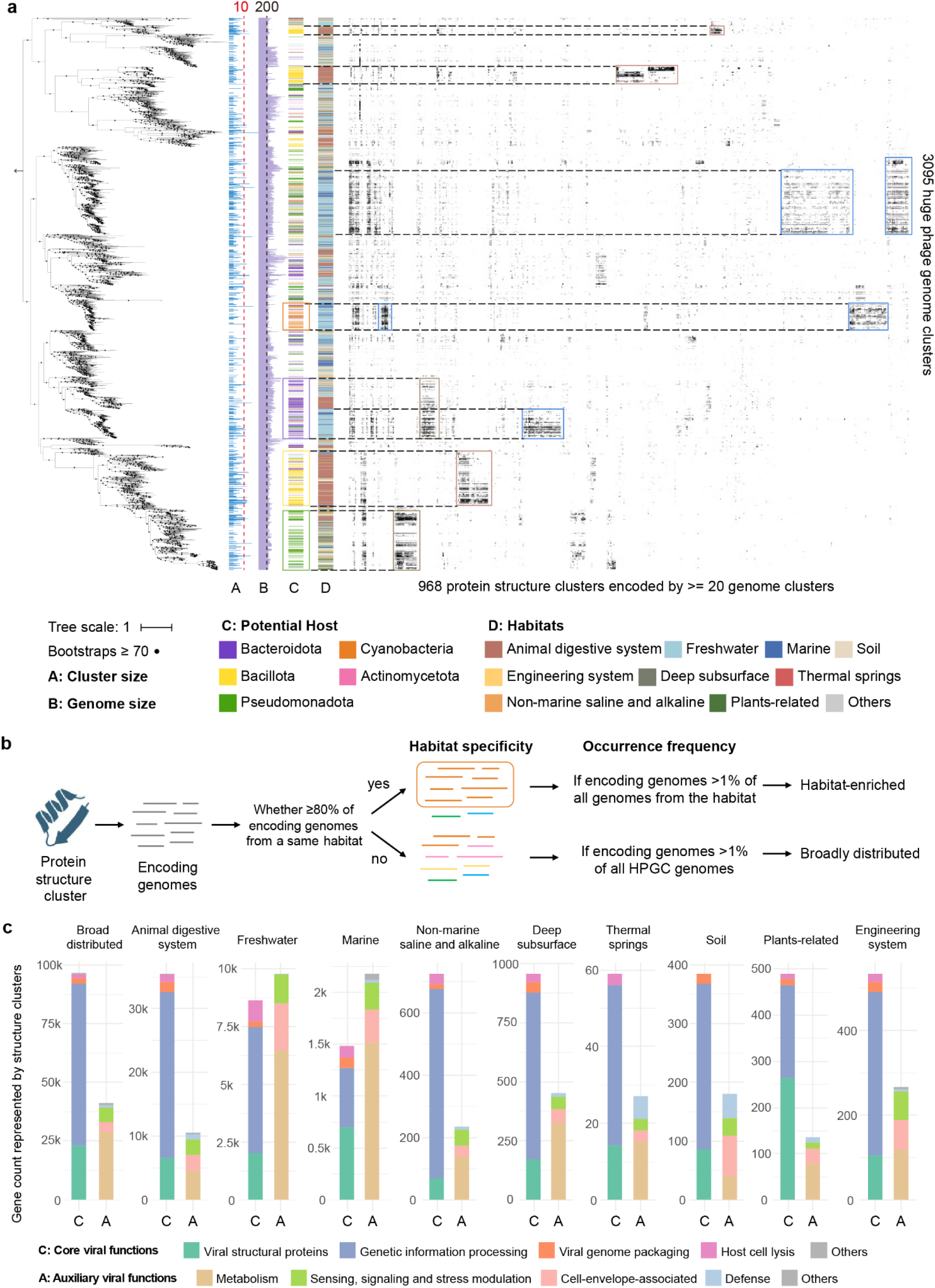
The distribution patterns of protein structure clusters among huge phages across diverse habitats. (a) TerL phylogenetic tree of huge phage genome clusters coupled with a presence–absence heatmap of protein structure clusters. Each leaf on the tree represents a huge phage genome cluster. The tree was rooted at the NCLDV A32 packaging proteins (indicated by an arrow to the left). Four annotation tracks surrounding the TerL tree represent: genome cluster size (log-transformed; reference line at cluster size = 10), genome length (reference line at 200 kb), predicted host phylum, and habitat origin. Rows in the heatmap follow the order of the TerL tree, and columns were ordered by hierarchical clustering based on Euclidean distances (Methods). Only protein clusters detected in at least 20 huge phage genome clusters are shown. In the heatmap, dark red, blue, and brown boxes indicate modular protein organization in the animal digestive system, aquatic habitats (including freshwater, marine, non-marine saline and alkaline, and deep subsurface), and multiple habitats, respectively. (b) The definition of habitat-enriched and broadly distributed protein structure clusters. A protein structure cluster is defined as habitat-enriched if ≥ 80% of the genomes encoding that protein originate from the same habitat, and it is present in >1% of genomes within that habitat. The remaining clusters that were detected in >1% of all HPGC genomes and found in multiple ecosystems were classified as broadly distributed. (c) Functional classification of habitat-enriched and broadly distributed protein structure clusters across ecosystems. Only protein clusters with available structure-based functional annotations were included and were assigned to core viral functions or auxiliary viral functions. Stacked bars indicate the total number of genes represented by protein structure clusters assigned to each functional category.

To further capture habitat-associated functional proteins, we defined a subset of HPSDB protein structure clusters as habitat-enriched or broadly distributed (Figure 2b). A protein structure cluster is considered enriched in a given habitat if ≥ 80% of the genomes encoding that protein originate from the corresponding habitat, and it is present in >1% of the total genomes of that habitat. The remaining protein structure clusters that were detected in >1% of all HPGC genomes (i.e., encoded by ≥72 genomes) and found in multiple ecosystems were classified as broadly distributed. A total of 6,401 and 708 protein structure clusters representing 176,979 and 167,040 proteins were categorized into habitat-enriched and broadly distributed groups, respectively (Supplementary Table 2).

The annotatable proteins within these two groups were subsequently assigned to either core viral functions or auxiliary viral functions (Figure 2c, Supplementary Table 2). Metabolism was the most frequently detected category of auxiliary viral functions, with protein classes such as glycosyltransferases and peptidases frequently represented among the broadly distributed group. Broadly distributed clusters also included PfkB-family carbohydrate kinases and glutamine synthetases, associated with carbohydrate metabolism ^42^ and nitrogen assimilation ^43^, respectively. Both proteins were predominantly detected in freshwater and marine huge phages. We speculate that the phage-encoded glutamine synthetase could enable them to directly modulate host nitrogen metabolism by synthesizing glutamine, a critical nitrogen carrier and precursor for nucleotides and amino acids, thus enhancing their fitness under nitrogen-limited conditions ^44^. The sulfatase maturation enzyme AslB, which is involved in organosulfur metabolism through sulfatase maturation ^45^, was found to be enriched in the animal digestive system, suggesting potential viral support for host utilization of sulfated substrates. Proteins involved in sensing, signaling, and stress modulation, cell envelope-associated functions, and defense were also detected in all ecosystems. These auxiliary viral proteins represent an accessory functional repertoire of huge phages, providing host manipulation strategies and potential ecological benefits that extend beyond essential viral propagation.

### Diverse cell envelope-associated proteins are encoded by huge phages from the human gut

Functional annotation revealed that huge phages encode diverse auxiliary functions beyond those directly involved in viral replication and assembly (Figure 2c). Among these, proteins related to the bacterial cell envelope were particularly notable because the cell envelope serves as a critical interface for solute transport, maintaining cellular homeostasis, and supporting interactions with the surrounding environment ^14,46^. Many of these cell envelope-associated proteins were detected in gut-associated huge phages, prompting us to investigate their diversity in detail.

To characterize cell envelope functions shared across multiple genomes, we selected protein structure clusters that were detected in at least 10 gut-derived huge phage genomes and had informative structure-based annotations. Functional screening identified 57 cell envelope-associated protein clusters, collectively representing 2,916 proteins (Figure 3a, Supplementary Table 3). Among these proteins, membrane transport-associated proteins formed the most diverse category and accounted for the largest number of proteins (1429, 42.1%), encompassing both broad transporter families (e.g., ATP-binding cassette, major facilitator superfamily, and drug/metabolite transporters) and transporters of specific metabolites and ions (e.g., nicotinamide riboside, formate/nitrite, calcium, and magnesium transporters) (Figure 3b).

**Figure 3.**
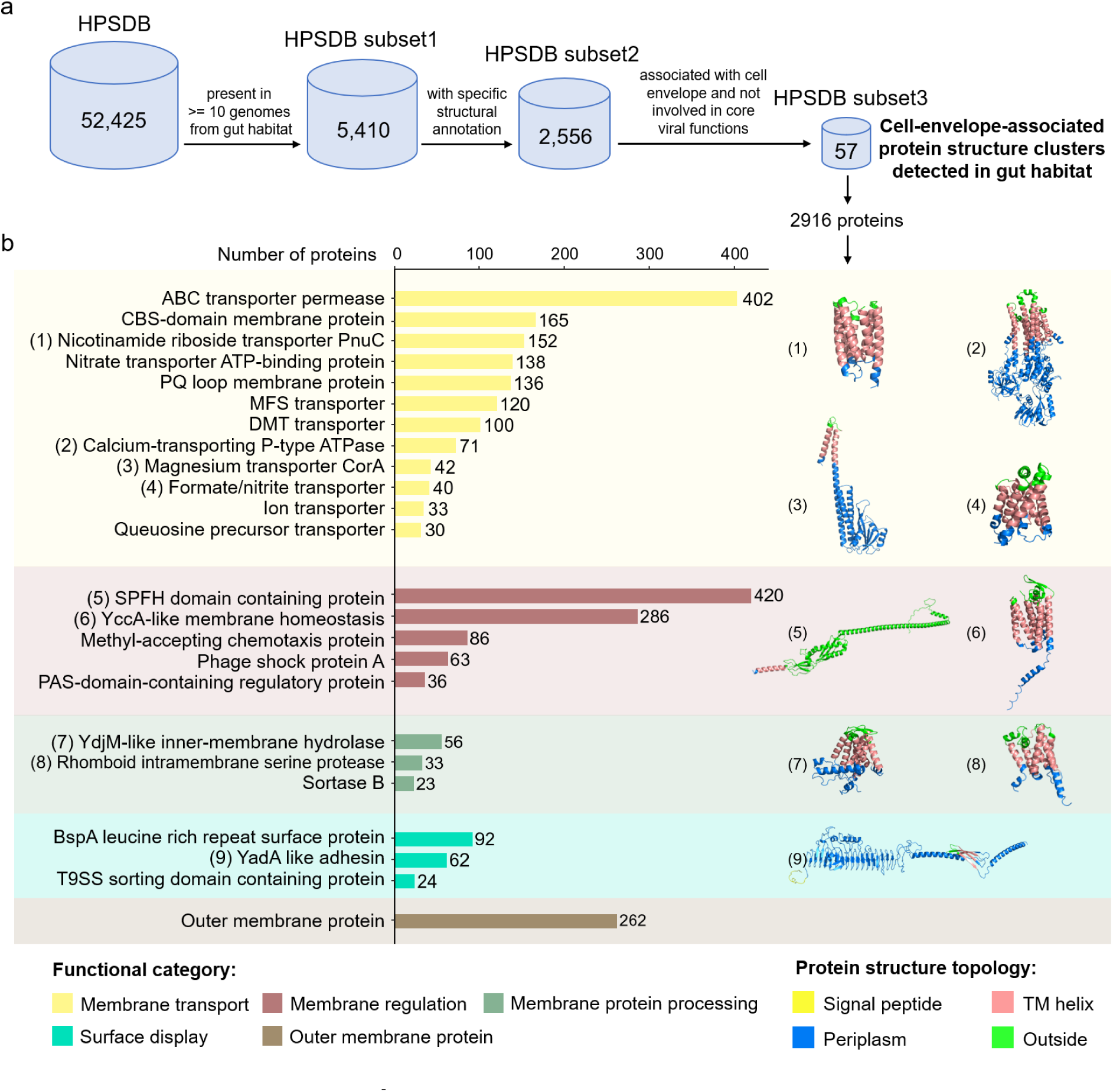
The host cell envelope-associated proteins encoded by gut-retrieved huge phages. (a) Screening workflow for identifying host cell envelope-associated protein structure clusters encoded by gut-retrieved huge phage genomes. Protein structure clusters presented in at least 10 huge phage genomes and with informative structure-based annotations were screened for cell envelope-associated functions, excluding those involved in core viral functions. The numbers indicate the number of protein structure clusters retained at each screening step. (b) Functional classification of host cell envelope-associated proteins. Horizontal bars indicate the total number of proteins assigned to the corresponding function. Examples of predicted structures of protein cluster representatives containing transmembrane regions are shown. The predicted structure topologies are indicated by different colors. The structures were predicted using ColabFold (Methods).

The next largest category comprised proteins associated with membrane regulation (891 proteins in total; Figure 3b). Among them, SPFH-domain containing proteins, which are associated with bacterial membrane organization and stress adaptation ^47,48^, were abundant (420 proteins). Proteins involved in membrane protein processing (112 proteins) and surface display (178 proteins) were also identified. Outer membrane proteins (OMPs; 262 proteins in total) were assigned to a separate category owing to their diverse functions (see details below). This functional overview highlights the diversity of cell envelope-associated proteins in gut-derived huge phages and provides the basis for further analysis of specific subsets in the following sections.

### Sortase B and substrate modules may mediate host cell-wall anchoring in gut-associated huge phages

We identified 24 sortase B (SrtB) genes encoded by 24 huge phages from the animal digestive system (Supplementary Table 4). Sortases, found primarily in Gram-positive bacteria, are membrane-associated cysteine transpeptidases that could covalently anchor their substrates to the cell-wall peptidoglycan ^49^. Their substrates are typically translocated across the cytoplasmic membrane via the Sec pathway, after which sortases recognize and cleave specific C-terminal cell-wall sorting signals and covalently attach the substrates to peptidoglycan ^50,51^. To our knowledge, sortase has not been documented in viral genomes but has occasionally been reported in other types of mobile genetic elements (MGEs), such as plasmids ^36,52^, extracellular chromosome elements ^53^, and integrative conjugative elements ^54^. Most sortase representatives identified in the non-viral MGEs belong to SrtA, while all those found in gut-associated huge phage genomes are SrtB (Figure 4a, left). Phylogenetic analysis showed that the SrtB encoded by huge phages were most closely related to those from Bacillota (Figure 4a, right).

**Figure 4.**
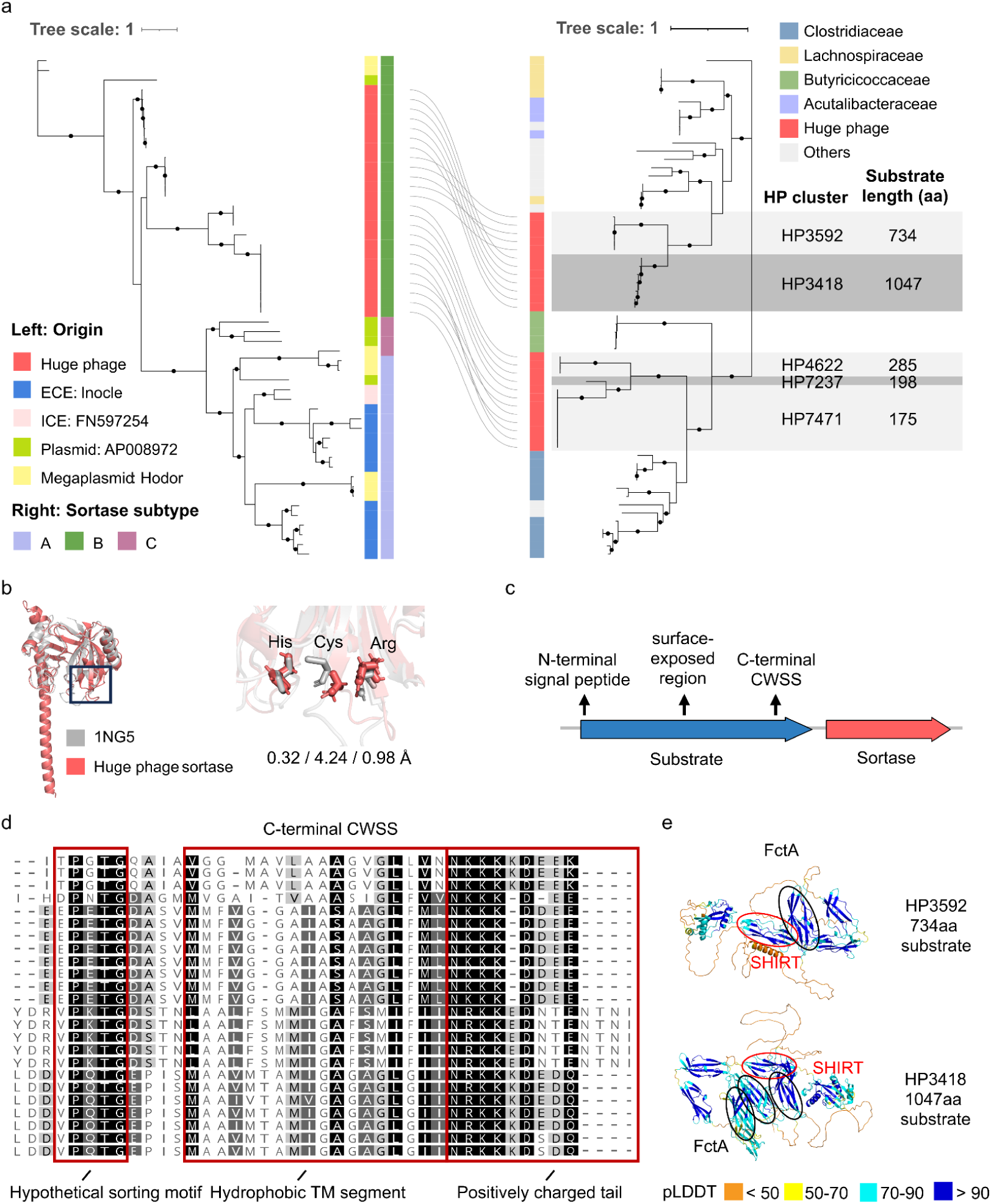
Phylogenetic analysis and structural features of sortases and their potential substrates in gut-associated huge phages. (a) Phylogenetic trees of sortases. Left: tree of sortases from this study and sortase representatives from other non-viral MGEs. Sortases from other MGEs were collected from four published references ^36,52–54^ and clustered with CD-HIT (parameters: -c 0.9 -aL 0.8 -aS 0.8), and only representative sequences were used in tree construction (Methods). Right: tree of sortases from this study and bacterial sortase reference sequences from the HRGM2 (Human Reference Gut Microbiome v2) protein database ^60^. The corresponding huge phage genome cluster and the length of the upstream potential substrate of sortase were indicated by a gray background. Solid black lines linking the two trees connect identical sortase sequences. (b) Structural comparison between huge phage SrtB (from huge_phage_2611 as an example) and the experimentally resolved bacterial SrtB (PDB ID: 1NG5). The right panel zooms in on the catalytic residues (i.e., His, Cys, and Arg, which are included in the black box in the left panel), with positional deviations indicated. (c) Schematic figure showing the SrtB gene and its upstream potential substrate gene, with the primary substrate architecture annotated. (d) Truncated sequence alignment showing the C-terminal cell-wall sorting regions of potential substrates, with the conserved sorting motif, hydrophobic segment, and positively charged tail highlighted with red boxes. (e) Predicted structures of the potential SrtB substrates from the HP3592 and HP3418 genome clusters. The SHIRT domains are indicated by red circles, and the FctA domains by black circles. The pLDDT values are shown by different colors.

Sequence alignment suggested that all huge phage SrtB proteins retained the conserved His, Cys, and Arg residues associated with sortase catalysis (Supplementary Figure 2). Structural superposition of the huge phage SrtB structure onto the experimentally determined bacterial SrtB structure (PDB 1NG5) placed these three residues in closely corresponding positions, with deviations of 0.32 Å (His), 4.24 Å (Cys), and 0.98 Å (Arg), respectively (Figure 4b).

SrtB-anchored substrates can support diverse cell-surface functions, including nutrient acquisition and adhesion ^55^, with haem–iron acquisition representing the best-characterized role of this system ^56^. In all SrtB-encoding huge phage genomes, the gene immediately upstream of the SrtB gene was predicted to encode a putative sortase anchor protein. These proteins exhibited the characteristic architecture of cell-wall-anchored surface proteins ^57^, with an N-terminal signal peptide, a variable surface-exposed region, and a C-terminal cell-wall sorting signal (CWSS) containing a conserved sorting motif, a hydrophobic segment, and a positively charged tail (Figures 4c and d). Alignment of potential SrtB-anchored substrates revealed a conserved PXTG sorting motif (Figure 4d). Despite the conservation of C-terminal CWSS regions, the substrates encoded by different huge phage genome clusters varied substantially in overall length, ranging from 175 to 1,047 amino acids (Figure 4a). The mean pLDDT values of these substrate proteins were below 70, largely reflecting low-confidence interdomain regions, whereas several folded domains showed high prediction confidence (pLDDT >90) (Figure 4e).

The predicted structures of the two longer substrates, encoded by the HP3418 and HP3592 genome clusters, both contained recognizable SHIRT (Streptococcal High Identity Repeats in Tandem) and FctA-like domains (Figure 4e). SHIRT domains are known to form elongated stalk-like structures that extend functional domains away from the bacterial cell surface and modulate their accessibility to extracellular interaction partners ^58^. FctA-like domains, in contrast, are associated with the structurally stable backbone of Gram-positive bacterial pili, which contribute to cell adhesion and other surface interactions ^59^. The presence of both domain types in these phage-encoded substrates suggests a potential structural organization that combines extended surface projection with mechanical stability, potentially facilitating interactions with extracellular molecules or neighboring cells. Although the specific functions of these substrates remain unknown, their shared structural features and conserved C-terminal anchoring signals suggest that some huge phages may encode specialized cell-surface proteins capable of modifying the surface properties of infected bacteria. More broadly, the conservation of SrtB-associated anchoring machinery across these phages, together with substantial variation in substrate length and domain architecture, suggests a modular system through which different phage lineages may deploy distinct extracellular proteins on bacterial cell surfaces.

### PspA as a potential modular effector for host membrane stress management in huge phages

PspA is the major effector of the bacterial phage shock protein system and directly contributes to cytoplasmic membrane integrity and remodeling. Structural and functional studies have shown that PspA forms ESCRT-III (Endosomal Sorting Complex Required for Transport III)-like assemblies capable of interacting with lipids, inducing membrane curvature, and promoting membrane remodeling ^61^. A total of 189 *psp*A genes were identified in 184 huge phage genomes, five of which carried two *psp*A copies each. These genomes originated from diverse ecosystems, most of which were recovered from animal digestive systems (all from the human gut, n = 118), followed by soil (n = 34) and freshwater (n = 9) (Supplementary Table 5).

The phylogenetic analysis of PspA proteins included gut-derived reference sequences retrieved from the HRGM2 protein database ^60^, most of which were encoded on scaffolds belonging to Bacillota MAGs (Figure 5a). Viral screening of the HRGM2 *psp*A-encoding sequences revealed that more than half of these reference *psp*A genes were located on phage-related scaffolds within the bacterial MAGs. This high representation of *pspA* genes in phages motivated a broader examination, leading to the identification of 667 high-confidence phage genomes (length range: 8.8-548 kbp) from IMG/VR that also encoded *psp*A (Supplementary Table 6). Among these IMG/VR genomes, 345 had an estimated completeness of at least 90%, and their genome lengths ranged from 41 to 548 kbp (∼168 kbp on average), indicating that viral *psp*A genes were not rare across ecosystems and were generally encoded by those with relatively large genome sizes (Figure 5b).

**Figure 5.**
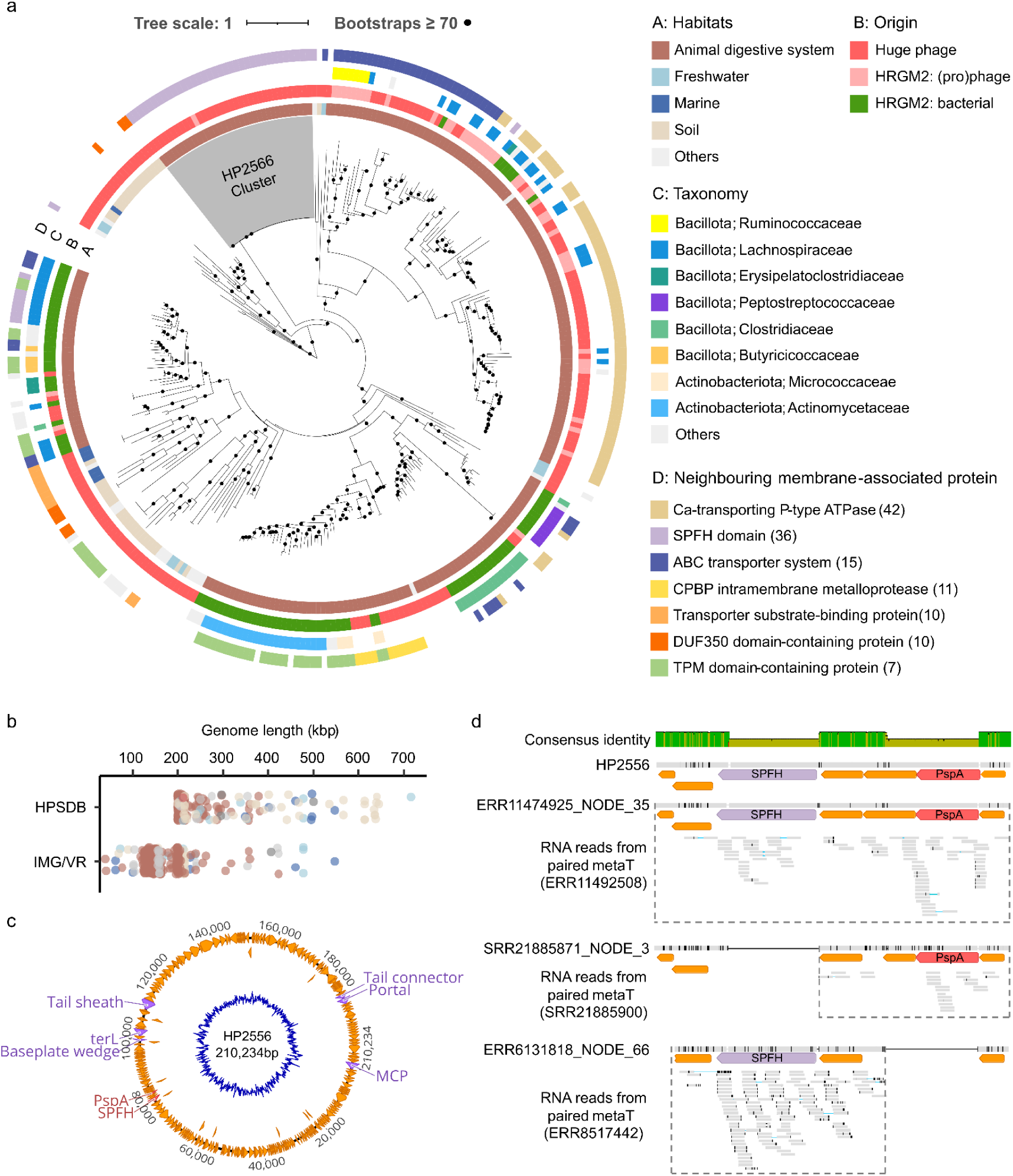
Phylogenetic distribution, genomic context, and transcriptional activity of huge phage-encoded PspA. (a) Phylogenetic tree of PspA sequences from this study together with reference PspA sequences from the HRGM2 protein database ^60^. Four color strips indicate habitat, sequence origin, taxonomic classification, and the type of neighboring membrane-associated protein. Taxonomy labels are shown only for HRGM2 reference sequences originating from bacterial or prophage PspA; full-length phage contigs binned into MAGs were assigned as unknown, thus the taxonomy was not shown on the tree, as the co-binning of full-length phage contigs into MAGs may not be reliable ^68^. Numbers next to the membrane-associated protein legend denote the detection count for each protein type. The grey colored range on the tree highlights PspA sequences from the HP2556 genome cluster. (b) Genome length distribution of PspA-encoding huge phages from this study and high-confidence vOTUs (and also with estimated completeness ≥90%) from IMG/VR version 7.1. Dot colors represent habitat categories, consistent with the legend in (a). (c) The complete huge phage genome HP2556 (210,234 bp in length), with selected viral marker genes, PspA and the SPFH-domain genes highlighted. (d) Alignment of the genomic region surrounding PspA in HP2556 and three related assembled genomes, with mapped RNA reads showing transcriptional activity.

Characterization of the genomic neighborhoods (the five upstream and five downstream genes) surrounding huge phage-encoded *psp*A genes revealed that 133 of the 189 loci (70.4%) contained other membrane-associated proteins (Supplementary Table 5). The neighborhood configurations formed distinct clades on the PspA phylogenetic tree, indicating evolutionarily structured gene arrangements (Figure 5a). The most frequently found membrane-associated protein near PspA was a Ca²⁺-transporting P-type ATPase (42 cases), which is expected to transport Ca²⁺ across the cytoplasmic membrane in bacteria ^62,63^ and has been reported in Jug phages detected in the gut microbiomes ^23^. The corresponding PspA proteins related to Ca²⁺-transporting P-type ATPases were most closely related to those from Lachnospiraceae within Bacillota. TerD and VWA-domain proteins, which were implicated in metal-dependent binding ^64,65^, were commonly encoded between PspA and the Ca²⁺-transporting ATPase, suggesting a putative cation-associated membrane-stress module (Figure S3).

The second prominent membrane-associated protein near PspA was an SPFH-domain protein (36 cases) (Supplementary Figure 3), which participates in membrane organization and stress adaptation in bacteria ^66^. In addition, complete ABC transporter systems were found adjacent to 15 PspA genes, each comprising a substrate-binding protein, an ATP-binding protein, and a transmembrane permease, supporting the capacity for ATP-dependent substrate transport and possible cooperation with PspA in membrane stress mitigation. Other membrane-associated proteins near PspA included a CPBP intramembrane metalloprotease (11 cases), a predicted two-transmembrane-helix transporter substrate-binding protein (10 cases), a DUF350 domain-containing protein (10 cases), and a TPM domain-containing protein (7 cases). These neighborhood configurations positioned PspA as a recurrent membrane-stress component embedded within diverse membrane-related modules.

To reveal the transcriptional activity of phage-encoded *pspA* genes, we used PebbleScout searching ^67^, with *pspA*-encoding huge phage genomes as queries (Methods). As a result, we identified HP2556-related huge phages in three NCBI Bioprojects with paired metagenomic and metatranscriptomic datasets, all of which were from the human gut. The reference genome HP2556 was reconstructed from the human gut with a circular genome supported by an identical terminal overlap (Figure 5c). HP2556 encoded an SPFH-domain protein separated from PspA by two hypothetical proteins. We performed *de novo* assembly on the Pebblescout-retrieved metagenomic samples and yielded three HP2556-related phage genomes. Whole-genome comparison showed that all three assembled genomes were highly similar to HP2556, with average nucleotide identities of 96.91–99.78% and query alignment fractions of 95.59–97.06% (Supplementary Figure 4). Alignment of the regions surrounding PspA in the three Pebblescout genomes against HP2556 revealed that the genome recovered from ERR11474925 also encoded both PspA and the SPFH-domain protein, while the related genomes from SRR21885871 and ERR6131818 encoded only PspA and only the SPFH-domain protein, respectively (Figure 5d). This patchy distribution is consistent with differential gain or loss of these membrane-associated functions and indicates that PspA is not restricted to a fixed genomic association with the SPFH-domain protein. Metatranscriptomic reads further supported the expression of the retained genes in their respective genomes, demonstrating that these evolutionarily dynamic membrane-associated functions are transcriptionally active in natural microbial communities.

### Various types of outer membrane proteins encoded by huge phages from the gut and other habitats

We also identified OMP proteins with β-barrel domains encoded by huge phages from the human gut and other ecosystems (Figure 6a). OMPs are integral components of the outer membrane in Gram-negative bacteria and perform diverse functions in envelope stabilization, solute transport, surface display, and extracellular interactions ^69^. Notably, bacterial OMPs with β-barrels ranging from 8 to 22 transmembrane strands have been experimentally shown to serve as receptors for phage infection ^70–73^. By integrating annotations with transmembrane topology analyses (Methods), 1,603 putative huge phage-encoded OMPs with predicted β-barrels containing 6–22 transmembrane strands were identified, including 678 from freshwater, 297 from animal digestive systems, 296 from marine environments, and 189 from soil (Supplementary Table 7). These OMPs were identified in 1,303 huge phage genomes (18.1% of HPGC), 250 of which encoded two or more OMPs (up to 4). The composition of the huge phage-encoded OMPs repertoire varied markedly among ecosystems. OMPs from animal digestive systems were dominated by 8- and 22-stranded β-barrels, while those from soil, freshwater and marine environments encompassed a broader range of barrel sizes.

**Figure 6.**
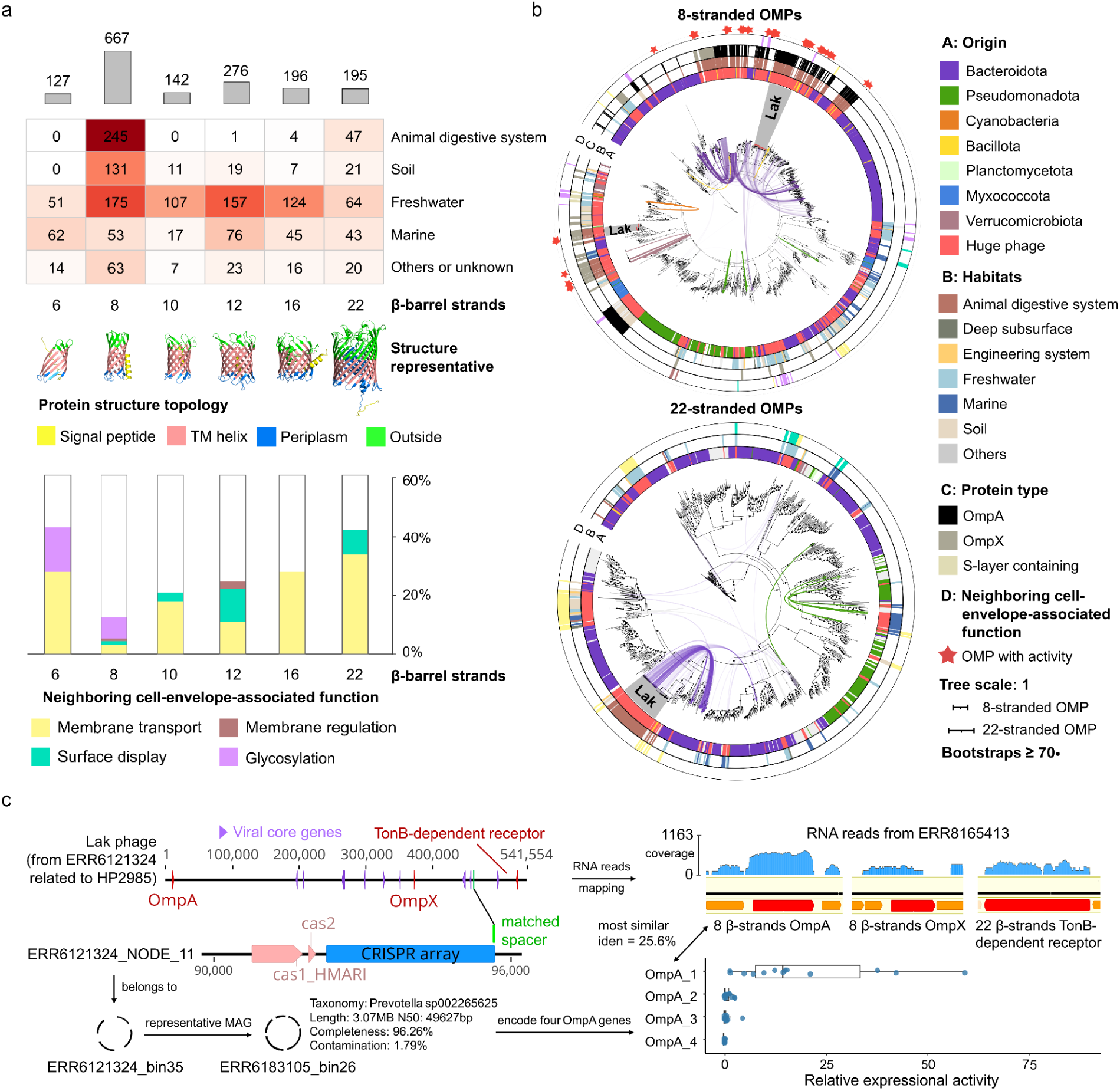
The diversity, phylogeny, and transcriptional activity of OMPs in huge phages. (a) Numbers of OMPs with 6–22 predicted β-barrel strands across ecosystems (top), representative structures (middle), and encoding patterns of genomic neighborhoods of OMPs associated with the cell envelope (bottom). Glycosyltransferases were functionally assigned to AMGs in this study and were also counted as cell envelope-associated genes here because they can be involved in envelope glycan biosynthesis. (b) Phylogenetic trees of 8-stranded and 22-stranded OMPs. The trees comprise huge phage OMPs identified in this study, OMP homologs retrieved from NCBI nr, and homologs encoded by bacterial genomes predicted as huge phage hosts using iPHoP. Curves connect huge phage OMPs to homologs in their predicted host genomes. Red stars mark 8-stranded OMPs assembled from public datasets for which transcription was detected. Outer color strips indicate taxonomic origin (A), habitat (B), protein type (C, only for 8-stranded OMPs), and neighboring gene function (D), respectively. Color strip D is the same as those in (a). (c) Transcriptional activity of OMP genes encoded by a Lak phage MAG and its predicted host MAG. The left panel shows a Lak phage genome encoding genes for OmpA, OmpX, and a TonB-dependent receptor. The genome was assembled from ERR6121324 and is related to the HP2985 genome cluster. CRISPR spacer match linked this phage to a host MAG assigned to *Prevotella* sp002265625, which encodes four *ompA* genes. The right panel shows the transcriptional activity of the three phage OMP genes and the four host *ompA* genes. Expression of the phage OMP genes was quantified by mapping reads from the paired metatranscriptome ERR8165413 to the Lak phage genome. Expression of the host *ompA* genes was quantified by mapping reads from 14 metatranscriptomes from BioProject PRJEB45356 to the host MAG (Methods). The host OmpA homolog most similar to phage OmpA (25.6% overall amino acid identity), i.e., OmpA_1, showed the highest transcriptional activity among the four host OmpA homologs.

Almost all bacterial homologs of the gut-derived OMPs, as well as the predicted hosts of the corresponding huge phages, were assigned to Bacteroidota, with the exception of a minor lineage of gut-derived 8-stranded OMPs associated with Verrucomicrobiota (Figure 6b). In contrast, huge phage OMPs from other ecosystems were more frequently associated with Pseudomonadota and Cyanobacteriota. We also identified a distinct 16-stranded OMP lineage whose phage hosts were assigned to Selenomonadaceae, a group that is exceptional among Bacillota for possessing an outer membrane (Supplementary Figure 5).

Most identified 6-stranded OMPs were encoded by huge phages from aquatic environments (Supplementary Table 7) and were related to the previously described cyanophage-encoded OMP-like family IPR056410 (also known as DUF680) ^74^. The putative membrane β-barrel of IPR056410 is incomplete, but structural modeling predicts that two subunits can assemble into a flattened 12-stranded β-barrel (Supplementary Figure 5). These OMPs occurred in different genomic contexts, including phosphate-acquisition modules and conserved virion-assembly loci. The 8-stranded β-barrels OMPs constituted the most abundant class and represented multiple phylogenetically distinct protein types, including OmpA, OmpX, and S-layer-associated proteins (Figure 6b). OmpA primarily contributes to envelope stabilization ^75^, while OmpX and S-layer-associated proteins are associated with extracellular interactions ^76^ and cell-surface organization ^77^, respectively. Notably, cell envelope-associated genes were frequently identified in the genomic neighborhoods of 10-, 12-, 16-, and 22-stranded OMPs. Most of these associated genes are transport-related, such as genes for phosphate acquisition in phages from aquatic environments, and those for nitrate, ammonium, and nicotinamide mononucleotide transport were also detected (Figures 6a and b, Supplementary Table 7). However, transport- and other cell envelope-related genes were rarely found adjacent to 8-stranded OMPs, which typically do not form large solute-transport channels ^78^.

To investigate the evolutionary relationships between huge phage OMPs and host-encoded homologs, we included OMP homologs from putative host genomes for phylogenetic analysis (Figure 6b, Supplementary Figure S5). Phage OMPs were typically more closely related to homologs from their predicted hosts than to those from other phages and bacteria. This pattern is consistent with a possible history of OMP gene exchange between huge phages and their bacterial host lineages. In a few cases, phage OMPs showed relatively distant relationships to the recovered host homologs. This suggests cross-lineage OMP gene exchange, or alternatively, the corresponding bacteria may encode the gene exchange source of the phage-encoded OMP genes, but the scaffolds were not binned into the MAGs.

Lak phage represents a group of huge phages widely detected in human and animal gut microbiomes and could infect *Prevotella* spp. ^20,79^. Notably, some Lak phages belong to the HP2985 genome cluster, which simultaneously encodes an 8-stranded OmpA protein, an 8-stranded OmpX-like protein, and a 22-stranded TonB-dependent receptor, all of which displayed detectable transcriptional activity (Figure 6c). CRISPR spacer matches and iPHoP predictions linked this phage to a putative host MAG assigned to *Prevotella* sp002265625, which encoded four *ompA* genes. Among the four host *ompA* genes, OmpA_1 consistently showed the highest transcriptional activity in all 14 samples (Figure 6c, Supplementary Table 8). The OmpA_1 protein showed the closest match to HP2985 OmpA at the sequence level (25.6% amino acid identity) and the structural level (DALI Z-score = 19.4, RMSD = 4.7 Å over 210 aligned residues) (Supplementary Table 9).

We then comprehensively examined the expression of 8-stranded OMPs encoded by gut-associated huge phages using publicly available metatranscriptomic datasets and detected their transcriptional activity across several major phylogenetic clades (Figure 6b). These 8-stranded OMP genes were generally expressed at significantly higher levels than the median of other active genes encoded by the same phages, and in some cases ranked among the top 10 most highly expressed genes (Supplementary Table 10). Transcriptional evidence for OMPs with other β-barrel sizes was also observed (see examples in Supplementary Figure 6).

## Discussion

The rapid expansion of metagenomic sequencing and development of viral analysis tools ^80–82^ have identified thousands of huge phage genomes ^17,23^, yet their functional diversity remains difficult to resolve because a large fraction of their proteins show limited sequence similarity to experimentally characterized proteins ^24^. HPSDB provides a complementary view of these poorly characterized proteins by analyzing the dominant proteins encoded by 7,187 non-redundant huge phages according to their predicted structures (Figure 1). The majority of the HPSDB structure clusters were represented by a single predicted structure, suggesting that the expanded genomes of huge phages encode a remarkably diverse protein repertoire. This was further supported as more than half of the structural cluster representatives lacked significant matches to the two published virus-derived structural databases ^37,38^ (Figure 1d). HPSDB therefore serves not only as an annotation resource but also as a framework for prioritizing unexplored proteins according to their structural conservation, phylogenetic distribution, ecological association, and genomic context (Figure 2).

The huge phage protein repertoires appear to comprise a relatively conserved component and a much more diverse accessory repertoire. Mapping abundant structural clusters onto the TerL phylogeny showed that related phages frequently shared characteristic combinations of structurally similar proteins (Figure 2a), consistent with the inheritance of proteins involved in genome replication, virion assembly, and other core viral processes along with the viral genome. A similar pattern emerged from their ecological distributions. Although substantially fewer structural clusters were classified as broadly distributed than as habitat-enriched, the two groups represented similar numbers of proteins (Figure 2c), indicating that broadly distributed clusters comprise, on average, much larger protein families. Together, these patterns are consistent with a set of recurrent and relatively conserved functions shared across diverse huge phages and ecological settings. In contrast, the accessory repertoire appears substantially more diverse and ecologically differentiated. Numerous structural clusters showed restricted phylogenetic distributions or enrichment in particular habitats, and many included auxiliary functions related to metabolism, sensing and stress responses, defense, and host cell envelope (Figure 2c, Supplementary Table 2). Such variation may reflect multiple selective contexts. Phages infecting phylogenetically distinct bacteria encounter different cellular architectures, metabolic capacities, defense systems, and physiological constraints ^3,83,84^, potentially favoring different accessory functions, while environmental conditions may further influence their value by altering nutrient availability, physicochemical stress, and host physiological state. Consistent with this interpretation, variation in protein repertoires corresponded with both predicted host taxonomy and environmental origin (Figure 2). These associations cannot be readily disentangled from shared phage evolutionary history or from correlations between host distribution and habitat, but collectively suggest that the accessory repertoire provides a more flexible component of huge-phage genomes through which functional diversification can occur across different host and ecological contexts.

Large-genome phages possess expanded coding capacities that may enable them to engage with a broader range of host cellular processes ^17,21,24^. Previous studies have illustrated the diverse strategies by which large phages can modify intracellular processes ^6,25^. Our findings extend this view to the bacterial cell envelope, suggesting that the expanded accessory repertoires of large phages may enable them to modulate diverse aspects of host cell-envelope physiology, potentially through distinct molecular mechanisms.

For phage-encoded OMPs, one intriguing possibility is that some of them modify the receptor landscape of infected cells and thereby influence subsequent phage adsorption. This possibility is supported by a recent study of phage λ, which demonstrated that the phage-encoded 8-stranded β-barrel outer membrane protein Lom binds to the host receptor LamB and blocks its recognition by incoming λ phages, thereby preventing superinfection ^85^. The diverse β-barrel OMPs identified here in huge phages (Figure 6a) raise the possibility that similar receptor-level interference mechanisms may be more widespread than previously recognized. Beyond specific receptor binding, phage-encoded OMPs might also influence receptor accessibility through other mechanisms. For example, their expression during infection could potentially affect the assembly or spatial organization of host OMPs, or sterically interfere with the accessibility of endogenous surface receptors. Such changes could selectively reduce the adsorption of incoming phages that depend on particular receptors, while leaving other receptor pathways accessible. In this way, remodeling of the outer membrane protein landscape could provide a selective form of superinfection exclusion by reducing the adsorption of incoming phages that depend on particular host receptors. This possibility is particularly relevant to the transcriptionally active OMPs identified in this study (Figure 6c, Supplementary Table 10), although transcriptional activity alone does not establish protein abundance or function. Determining whether these proteins contribute to receptor interference or superinfection exclusion will require experimental validation of their outer membrane localization, interaction partners, and effects on phage adsorption.

Surface display through the SrtB and substrate systems could add yet another dimension to this remodeling of infected cells. SrtB could covalently expose phage-encoded proteins on the bacterial cell surface. The combination of a conserved SrtB-mediated anchoring mechanism with structurally diverse predicted substrates suggests that this system could display a variety of phage-encoded proteins on the bacterial cell surface (Figure 4). The variable extracellular regions could mediate diverse interactions at the bacterial cell surface and potentially alter the accessibility or organization of existing host surface structures. It is therefore conceivable that surface display contributes to receptor masking or other forms of superinfection interference, although this possibility remains more speculative than the receptor-level mechanisms proposed for OMPs.

Phage-encoded PspA suggests a different but potentially complementary strategy. Bacterial PspA responds to membrane stress, associates with phospholipids, and can reduce proton leakage from damaged membranes, thereby contributing to membrane integrity and maintenance of the proton motive force ^86^. Large-phage replication and virion assembly are likely to impose substantial energetic and membrane-associated stresses on infected cells ^87,88^. Expression of phage-encoded PspA could therefore help stabilize the cytoplasmic membrane or preserve membrane energetics during the productive phase of infection, effectively prolonging the physiological competence of the host until virion assembly is complete. Its recurrent association with Ca²⁺ P-type ATPases, SPFH-domain proteins, transporters, and other membrane-associated functions further raises the possibility that PspA operates as part of broader phage-directed membrane-management programs rather than as an isolated stress-response protein (Figure 5a). The observed transcriptional activity of representative PspA-associated loci (Figure 5d) is consistent with such a role, although the timing of expression during infection and its direct consequences for membrane physiology remain to be determined.

More generally, the three systems examined here illustrate how phage-encoded proteins could act at different layers of the bacterial envelope, from cytoplasmic membrane maintenance to outer-membrane remodeling and extracellular surface display. Together, these observations suggest that large-genome phages may engage with the bacterial cell envelope in two complementary ways: by maintaining or redirecting envelope physiology and by remodeling the interactions of infected cells with their extracellular environment. By integrating structural genomics with evolutionary, ecological, and gene transcription information, our study expands the view of the bacterial cell envelope from a barrier encountered during infection to a diverse physiological system potentially influenced by phage-encoded accessory proteins. Experimental characterization of these proteins will be essential to determine their biological functions and to better understand how expanded coding capacities contribute to the complex interactions between large-genome phages and their bacterial hosts.

## Methods

### Construction of the Huge Phage Protein Structure DataBase (HPSDB)

The Huge Phage protein structure DataBase (HPSDB) was constructed based on the Huge Phage Genome Collection (HPGC) ^23^. HPGC initially reported 7,230 huge phage genomes originating from diverse habitats, and 43 of them were further identified as non-viral genomes (some were megaplasmids ^36^). The remaining 7,187 huge phage genomes were subsequently clustered into 4,223 genome clusters using the scripts provided by the CheckV toolkit (https://bitbucket.org/berkeleylab/checkv/src/master/) with ≥90% average nucleotide identity across ≥80% alignment fraction ^80^.

Prodigal version 2.6.3 (single mode) ^89^ was used to predict protein-coding genes for each genome using genetic codes 4, 11, and 15, as previously described ^90^. In brief, the coding density of each genome was calculated by adding the total number of coding bases and dividing by the total genome length. The genomes with a coding density increase of ≥5% in code 4 or code 15 relative to code 11 were assigned that genetic code.

The 2,500,924 predicted proteins with the applied genetic code were then clustered using CD-HIT version 4.8.1 ^91^ with the parameters “-c 0.9 -aL 0.8 -aS 0.8 -G 0”, resulting in 1,509,176 sequence clusters. Among them, 162,625 clusters had members detected in ≥3 huge phage genomes, and their representatives were extracted for protein structure prediction using ColabFold (downloaded on 22 Nov 2024) ^32^. The structure prediction was conducted with the following parameters: “--num-recycle 3, --use-gpu-relax, --amber, --stop-at-score 70”. The best (“rank_001”) model structures with an average pLDDT score of ≥70 were clustered using the FoldSeek version 9.427df8a ^35^ “easy-cluster” command with the parameters of “-c = 0.7, --cov-mode = 0, --tmscore-threshold = 0.7”. These steps finally produced 52,425 protein structure clusters (Figure 1a).

### Protein structure comparison and annotation

To evaluate the novelty of the HPSDB structures, we searched the 52,425 structure cluster representatives against the two virus-derived reference structure databases, i.e., the Big Fantastic Virus Database (BFVD) ^37^ and the eukaryotic viruses structure database ^38^, and also the PDB ^40^, and the initial release of the AlphaFold database that contains more than 500,000 proteins from eukaryotes, bacteria, and archaea without viruses (AFDB non-viral) ^92^.

For comprehensive functional annotation of HPSDB structures, we further searched cluster representatives against the AlphaFold Protein Structure Database clustered at 50% sequence identity (AFDB50) ^39^, and structure databases recently derived from functional protein resources ^93^, including the Clusters of Orthologous Groups (COG) database ^94^, the High-quality Automated and Manual Annotation of Proteins (HAMAP) database ^95^, the NCBIfam database ^96^, and (8) the Pfam database ^97^.

Both structure comparison and annotation analyses were conducted using FoldSeek version 9.427df8a ^35^ with the “easy-search” command (-c 0.7 --cov-mode 0), and only those with an alntmscore ≥ 0.5 and E-value < 10^−5^ were thought to be reliable matches. For the identification of viral marker proteins, habitat-associated proteins, and cell envelope-associated proteins, functional assignments were based primarily on annotations from AFDB50, BFVD, PDB, and Pfam. When annotations conflicted across these databases, precedence was given to AFDB50, followed by BFVD, PDB, and Pfam.

### Phylogenetic analysis of the terminase large subunit and major capsid protein

To obtain a more comprehensive sequence recovery for the terL and MCP genes, the recruitment of the TerL and MCP was first based on HPSDB annotations (see section: Protein structure comparison and annotation) and supplemented by an additional iterative search. Using TerL as an example, all members belonging to protein structure clusters annotated as TerL were pooled together to form an initial set of huge phage TerL sequences. Then, all proteins from the huge phage genomes that lacked a TerL annotation were searched using local BLASTp ^98^ against the initial TerL set. Hits with an E-value ≤ 10^−5^ underwent the same protein structure prediction and comparison pipeline as described above. Briefly, the protein structures of the BLASTp hits were predicted using ColabFold (downloaded on 22 Nov 2024) ^32^ and compared against the structural databases using FoldSeek version 9.427df8a ^35^. Proteins annotated as TerL through this additional search were also incorporated into the huge phage TerL dataset. The same approach was also applied to identify additional MCP sequences. A single representative TerL (or MCP) sequence was selected for each of the huge phage genome clusters for phylogenetic analysis as follows: preference was given to the TerL (or MCP) encoded by the representative of each genome cluster; if unavailable, the longest TerL (or MCP) sequence within the genome cluster was selected. These steps generated a final set of 3,095 TerL and 3,019 MCP sequences for the 4,223 huge phage genome clusters.

For phylogenetic analysis, the huge phage TerL sequences were analyzed alongside the A32 packaging proteins from nucleocytoplasmic large DNA viruses (NCLDVs), and the huge phage MCP sequences were analyzed alongside NCLDV MCPs. A32 and MCP protein sequences from NCLDVs were retrieved from the NCBI Virus database and clustered using CD-HIT version 4.8.1 ^91^ with parameters “-c 0.9 -aS 0.9 -d 100” to reduce redundancy, yielding 59 A32 and 311 MCP sequences as outgroups in phylogenetic analysis. Multiple sequence alignment was performed using MUSCLE version 5.3.linux64 ^99^. The alignments were trimmed with trimAl version 1.4.rev15 ^100^ using the parameter “-gt 0.2” to remove columns with insufficient information. Phylogenetic trees were constructed with IQ-TREE version 2.4.0 ^101^, with the best-fitting substitution model selected automatically using ModelFinder (-m MFP). The trees were rerooted between the phage proteins and the NCLDV outgroup proteins. The phylogenetic tree was visualized in iToL version 7.6 ^102^.

### Definition and functional classification of habitat-associated protein clusters

To examine the distribution and functions of abundant protein structure clusters across ecosystems, habitat-enriched and broadly distributed protein clusters were identified within HPSDB based on habitat specificity and occurrence frequency. A protein structure cluster was designated as habitat-enriched when a single habitat accounted for ≥80% of its members, and the protein cluster occurred in >1% of HPGC genomes from that habitat. The remaining protein clusters that did not meet the enrichment criteria but were present in ≥72 HPGC genomes (>1% of the full collection) across two or more ecosystems were classified as broadly distributed.

Protein clusters from both groups were then assigned to core viral functions or auxiliary viral functions according to their structure-based annotations. The core viral function categories comprised viral structural proteins, genetic information processing, viral genome packaging, and host cell lysis. All proteins involved in DNA or RNA processing or metabolism were included in the genetic information processing category. Clusters lacking functional annotations or with ambiguous annotations were assigned as unknown.

### Identification of cell envelope-associated proteins

To identify putative cell envelope-associated proteins encoded by gut-associated huge phages, we first selected protein structure clusters from HPSDB that were present in at least 10 gut-associated huge phage genomes. Representative sequences of protein clusters with functional annotations were then analyzed using DeepTMHMM version 1.0 ^103^ to predict their transmembrane topology. Proteins containing predicted transmembrane regions with assigned functions unrelated to core viral processes were considered putative host cell envelope-associated proteins.

For the three protein groups selected for detailed investigation (i.e., sortB, PspA, and OMP), we performed a comprehensive search across HPGC huge phage genomes reconstructed from all ecosystems to maximize the recovery of related sequences. All predicted proteins in HPGC genomes were searched against seed sequences from the initially identified protein structure clusters using BLASTp with an E-value threshold of 10^−5^. The resulting candidates were subjected to structure prediction and annotation using the same workflow as described in the previous section. All candidate sequences were also searched against the Pfam database ^97^ using hmmsearch version 3.4 ^104^ to confirm the functional assignment.

The OMP candidates were subjected to additional quality control. N-terminal signal peptides were predicted using DeepTMHMM version 1.0 ^103^. The number of transmembrane β-strands was independently predicted from amino acid sequences using DeepTMHMM and from predicted structures using PolarBearal3 ^105^. Only proteins containing an N-terminal signal peptide and showing concordant β-strand counts between the two methods were retained. The dimeric structure of the cyanophage-associated OMP with an incomplete six-stranded β-barrel was predicted using AlphaFold-Multimer ^106^.

### Phylogenetic analysis of cell-envelope associated proteins

For sortase, because all sortases from this study were identified in gut-derived huge phage genomes, homolog sequences were retrieved from the human gut-associated HRGM2 protein database ^60^ using BLASTp, with phage sortases as references. For each query, the top 20 hits with E-value ≤ 10^-^⁵ were retained. The candidate sortase homologs were further screened against Pfam profiles using hmmsearch version 3.4 ^104^. Only sequences whose best Pfam matches corresponded to PF04203.13 were retained as reference sequences for phylogenetic analysis. An additional phylogenetic tree was constructed to compare huge phage-encoded sortases with those encoded by non-viral MGEs. Reference sortase sequences from non-viral MGEs were collected from four published studies ^36,52–54^ and clustered within each study using CD-HIT with the parameters “-c 0.9 -aL 0.8 -aS 0.8”. The representative sequences of the clusters were retained for tree construction alongside the huge phage sortases.

For PspA, which was predominantly identified in gut-associated huge phages, homolog sequences were also retrieved from the HRGM2 protein database ^60^ through BLASTp, using the same screening criteria as for sortases. The candidate homologs were further screened against Pfam profiles using hmmsearch version 3.4 ^104^. Only sequences whose best Pfam matches corresponded to PF04012.12 for PspA were retained as reference sequences for phylogenetic analysis.

To determine the genomic origins of the reference sequences from the HRGM2 database, scaffolds encoding the sortase or pspA homologs were screened using the same viral identification pipeline applied during HPGC construction ^23^. All sortase references shown in Figure 4a were classified as bacterial, whereas some PspA references in Figure 5a were encoded within predicted prophages or on standalone phage contigs. For prophage-encoded homologs, the taxonomic labels of the corresponding MAGs were retained. Homologs encoded on standalone phage contigs binned into MAGs were assigned as “unknown” because the co-binning of full-length phage contigs into MAGs may not be reliable ^68^.

For OMPs, which were prevalent in huge phages across diverse ecosystems, homolog sequences were first obtained from the NCBI nr database using BLASTp, retaining the top 20 hits with E-value ≤ 10⁻⁵ for each query. To reduce sequence redundancy, candidate homologs were clustered using CD-HIT version 4.8.1 ^91^ (-c 0.9 -aL 0.8 -aS 0.8), and cluster representatives were retained for subsequent analysis. The bacterial hosts of OMP-encoding huge phages were predicted by iPHoP version 1.3.3 ^107^. Based on the host genome accessions reported by iPHoP, the corresponding genomes were downloaded from NCBI, followed by protein-coding gene prediction using Prodigal version 2.6.3 ^89^. The predicted proteins were searched against the corresponding huge phage OMPs using BLASTp to identify putative host-encoded homologs (E-value < 0.01). All candidate OMP homologs from the NCBI nr database and predicted host genomes were subjected to transmembrane-topology analysis using DeepTMHMM version 1.0 ^103^. Only homologs with the same predicted number of transmembrane β-strands as the corresponding huge phage OMP were retained as reference sequences for phylogenetic analysis. If a predicted host genome encoded multiple OMP homologs that met the above criteria, the homolog with the highest sequence similarity to the corresponding huge phage OMP was selected as the reference sequence. For the 8-stranded OMPs, additional OMP sequences reconstructed from public datasets with detectable transcriptional activity were also included.

The huge phage protein sequences and reference homologs were aligned using MUSCLE version 5.3.linux64 ^99^. Gap-rich positions were removed with trimAl version 1.4.rev15 ^100^ using the parameter “-gt 0.2”. Phylogenetic trees were constructed with IQ-TREE version 2.4.0 ^101^ with the best-fitting substitution model selected automatically using ModelFinder (-m MFP). The phylogenetic trees were visualized in iToL version 7.6 ^102^.

### Gene neighborhood analysis

To investigate the functional context of huge phage-encoded sortases, PspA and OMPs, proteins encoded by the five genes immediately upstream and downstream of each target gene (thus 10 genes in total) were retrieved for functional annotation and membrane topology prediction.

Potential sortase substrates were screened based on the characteristic architecture of sortase-anchored surface proteins. Neighboring proteins with signal peptides predicted by DeepTMHMM version 1.0 ^103^ were examined for a C-terminal sorting motif followed by a transmembrane helix and a positively charged tail. Putative sorting motifs were inferred from sequence conservation in alignments generated with MUSCLE version 5.3.linux64 ^99^. The approximate locations of putative transmembrane helices were estimated from hydrophobic amino acid stretches, and positively charged tails were assessed based on at least two consecutive basic residues (lysine or arginine) at the C-terminus ^108^.

For PspA and OMPs, the neighboring proteins were searched against the NCBI nr protein database using DIAMOND BLASTp (E-value ≤ 10⁻⁵) to identify those potentially associated with cell envelope functions. The sequence-based annotations were then confirmed via structure-based annotations. As HPSDB included only CD-HIT clusters detected in at least three genomes (see section: Construction of the Huge Phage Protein Structure Database), proteins not included in HPSDB were predicted using ColabFold (downloaded on 22 Nov 2024) ^32^ and annotated using Foldseek version 9.427df8a ^35^, following the same annotation procedure as described above. For proteins already represented in HPSDB, structure annotations were obtained directly from the corresponding protein structure clusters. Only cell envelope-associated proteins with structural models having pLDDT scores ≥70 and consistent sequence- and structure-based functional annotations were retained. If multiple types of cell envelope-associated genes were present in the neighborhood of a target PspA or OMP gene, the encoding pattern was counted only once and shown in Figure 6 and classified according to the nearest such gene. All identified cell envelope-associated genes in each genetic neighborhood are listed in Supplementary Tables 7 and 8.

### Sequence and structural comparison of huge phage and bacterial sortase B

Two experimentally characterized bacterial sortase B sequences were retrieved from reviewed UniProtKB entries (Q2FZE3 and Q8Y588) and aligned with huge phage sortase B sequences using MUSCLE version 5.3.linux64 ^99^. The alignment was inspected for conservation of the His, Cys, and Arg residues. The predicted structures of huge phage sortase B representatives were superimposed onto the crystal structure of Staphylococcus aureus sortase B (PDB ID: 1NG5) using the align command in PyMOL version 3.0.3 ^109^. Following superposition, distances between the Cα atoms of corresponding His, Cys, and Arg residues in the phage sortase B and bacterial structures were measured to assess their spatial correspondence.

### Identification of pspA-encoding viral genomes in the IMG/VR database

The analysis was restricted to uncultivated virus genomes (UViGs) classified as high confidence in IMG/VR version 2022-09-20_7.1 ^110^. Based on the expected length of PspA, proteins of 200–300 amino acids encoded by high-confidence UViGs were searched against the Pfam profile PF04012 (PspA_IM30) using hmmsearch version 3.4 ^104^. Hits with E-value ≤ 10⁻⁵ were retained as putative PspA candidates. These candidates were subsequently searched against the full Pfam database ^97^ using hmmsearch, and only those with PF04012 as their best hit were retained. The corresponding viral genomes were reassessed using the same viral identification pipeline applied for HPGC construction ^23^. Only genomes that passed this screening were retained.

### Metatranscriptomic profiling of cell envelope-associated proteins

To evaluate the transcriptional activity of the cell envelope-associated proteins, genome-cluster representatives encoding the target proteins were used as references to retrieve relevant meta-omics datasets in the NCBI SRA using PebbleScout ^67^. Paired metagenomic and metatranscriptomic samples containing signals of the target huge phages were downloaded, using minimum coverage thresholds of ≥30 for metagenomes and ≥10 for metatranscriptomes. Metagenomic reads were assembled with metaSPAdes version 3.15.2 ^111^ with parameter ‘-k 21,33,55,77,99,127’. Assembled scaffolds ≥10 kb in length were compared to the corresponding huge phage reference genomes using BLASTn. Scaffolds were retained if they aligned at ≥90% nucleotide identity across ≥80% of their length against the corresponding reference huge phage genomes. The retained scaffolds collectively constituted a metagenome-assembled genome (MAG) related to the corresponding huge phage reference genome. Prodigal version 2.6.3 ^89^ was used to predict genes on huge phage MAGs. Then, metatranscriptomic reads were mapped to the huge phage MAGs using Bowtie2 with default parameters ^112^. The transcriptional activity of predicted genes was calculated by dividing the total number of RNA bases mapped to the gene by its length. This measure enabled the comparison of the target cell envelope-associated protein genes with other genes encoded by the same huge phage MAG in the sample.

For samples showing detectable transcription of the target genes, metatranscriptomic read alignments were manually inspected in Geneious Prime Build 2025-03-24 ^113^ to screen for potential DNA contamination. BAM files were imported and examined alongside the predicted gene annotations. Samples were retained only if (1) most intergenic regions lacked mapped reads, (2) transcription was detected for only a subset of predicted genes, and (3) transcriptional activity varied markedly among expressed genes. Those samples with RNA reads mapped generally evenly across the genome, and/or mapped well across intergenic regions, were thought to be contaminated with DNA reads and thus excluded from further analyses.

### Host identification for the OMP-encoding Lak phage cluster HP2985

Lak phages are gut-associated huge phages linked to the bacterial genus *Prevotella* ^20^. PebbleScout searches identified Lak phage HP2985-related MAGs in several metagenomic samples from BioProject PRJEB45356 (see section: Metatranscriptomic profiling of cell envelope-associated proteins). These OMP-encoding HP2985-related MAGs were selected for further investigation of their host association and the potential relationships between phage- and host-encoded OMPs.

HP2985-related phage MAGs were recovered from 29 fecal metagenomes collected from 11 individuals, and each MAG has a total length >70% of the HP2985 reference genome length. The assembled scaffolds were binned using MetaBAT2 version 2:2.15 ^114^ with default parameters to recover MAGs. Genome completeness and contamination were assessed using CheckM2 version 0.1.3 ^115^, and MAGs with >50% completeness and <10% contamination were retained. These MAGs were dereplicated using dRep version 3.5.0 ^116^, and the dereplicated genomes were taxonomically classified using GTDB-Tk version 2.4.1 ^117^.

Host associations were inferred through CRISPR spacer matching and iPHoP prediction. CRISPR arrays were identified in the dereplicated bacterial MAGs using PILER-CR version 1.06 ^118^ with default parameters, and their spacers were extracted. Cas genes were identified by screening predicted proteins from array-containing scaffolds against a custom collection of Cas hidden Markov models derived primarily from TIGRFAM ^119^. Spacer matches were retained only if a cas gene was detected within 2 kb of the corresponding CRISPR array. Scaffolds from each HP2985-related phage MAG were searched against the spacers predicted from the same sample using BLASTn with an E-value cutoff of 10⁻⁴ and only one mismatch allowed across the whole spacer length. This analysis identified a putative host MAG representative taxonomically assigned to *Prevotella* sp002265625, with 96.26% completeness and 1.79% contamination. The host assignment was also predicted using iPHoP version 1.3.3 ^107^, which also supported *Prevotella* sp002265625 as the host species for HP2985.

### Identification, comparison, and transcriptional profiling of *ompA* encoded by host MAG of Lak phage HP2985

Protein-coding genes of the putative HP2985 host MAG were predicted using Prodigal version 2.6.3 ^89^. The predicted proteins were searched against the Pfam database ^97^ using hmmsearch version 3.4 ^104^, and those with the best Pfam match as OmpA (PF00691.20) were retained as candidates. Their transmembrane topologies were examined using DeepTMHMM version 1.0 ^103^. The OmpA candidates predicted to contain an eight-stranded transmembrane β-barrel were identified as OmpA proteins. Four *ompA* genes were identified in the host MAG.

The OmpA protein encoded by HP2985 was compared with the four OmpA proteins encoded by its host MAG at the sequence and structural levels. Sequence comparisons were performed using local BLASTp ^98^. Structures of the four host OmpA proteins were predicted using ColabFold (downloaded on 22 Nov 2024) ^32^ and aligned with the HP2985 OmpA structure using DALI version 5 ^120^.

Paired metatranscriptomes were available for 14 of the 29 metagenomes from which the HP2985-related phage MAGs were recovered. RNA reads were mapped to the host MAG using Bowtie2 with default parameters ^112^. The relative transcriptional activity was calculated for host genes by dividing the total number of RNA bases mapped to the gene by its length and normalizing by metatranscriptomic library size.

## Data availability

All the protein structures generated in this study are available at FigShare via 10.6084/m9.figshare.34746123.

## Acknowledgements

L-X.C. is supported by the Research Program of the University of Science and Technology of China. We thank the Supercomputing Center at the University of Science and Technology of China for its support of ColabFold analyses.

## Author contributions

L.Y. and L-X.C. designed this study, analyzed the data, drafted and finalized the manuscript, and approved the final version of the manuscript.

