## Supplementary Figures for "Large-genome phages encode diverse proteins associated with host cell envelope functions"

for

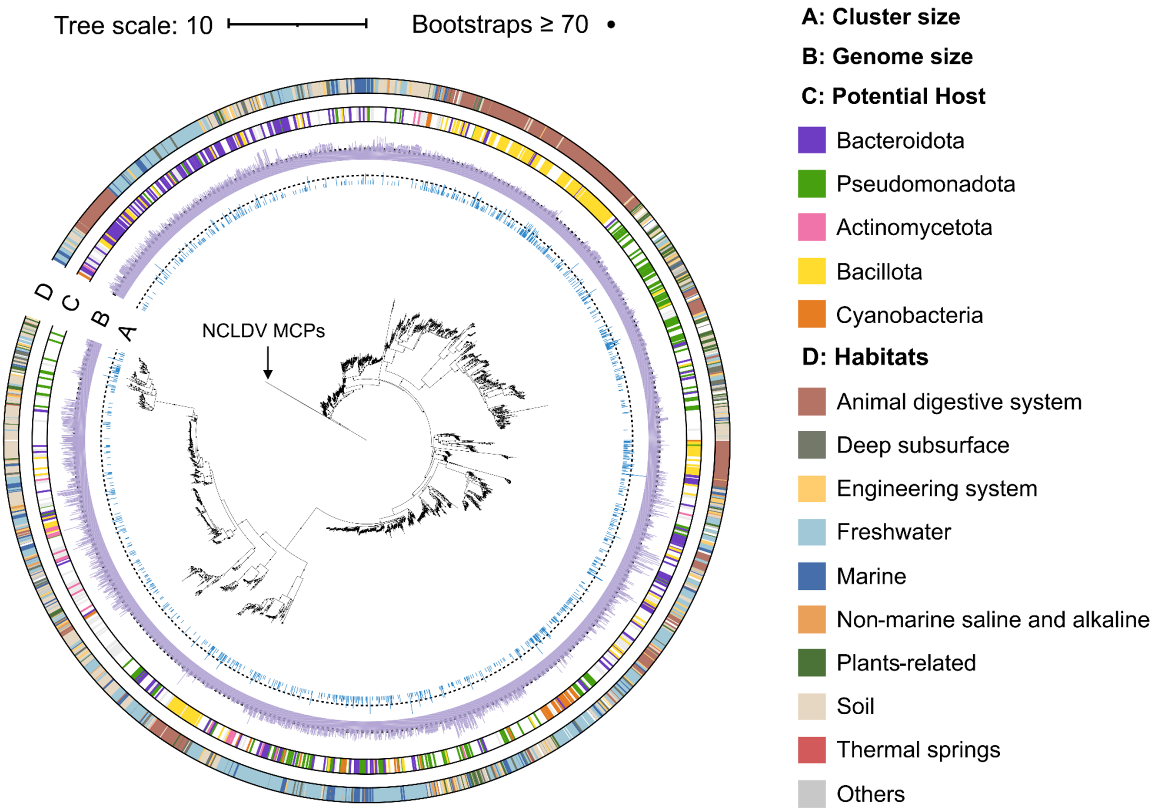


**Supplementary Figure S1 | The phylogeny of HPGC huge phages based on major capsid proteins.** Each leaf represents a genome cluster that was defined at at least 90 ANI across at least 80% of the genome length. Four annotation tracks surrounding the tree represent: genome cluster size (log-transformed; reference line at cluster size = 5), genome length (reference line at 200 kb), predicted host phylum, and habitat origin. The black dots indicate bootstraps, with only those ⩾70 shown. The tree was rooted between huge phage MCPs and NCLDV MCPs.

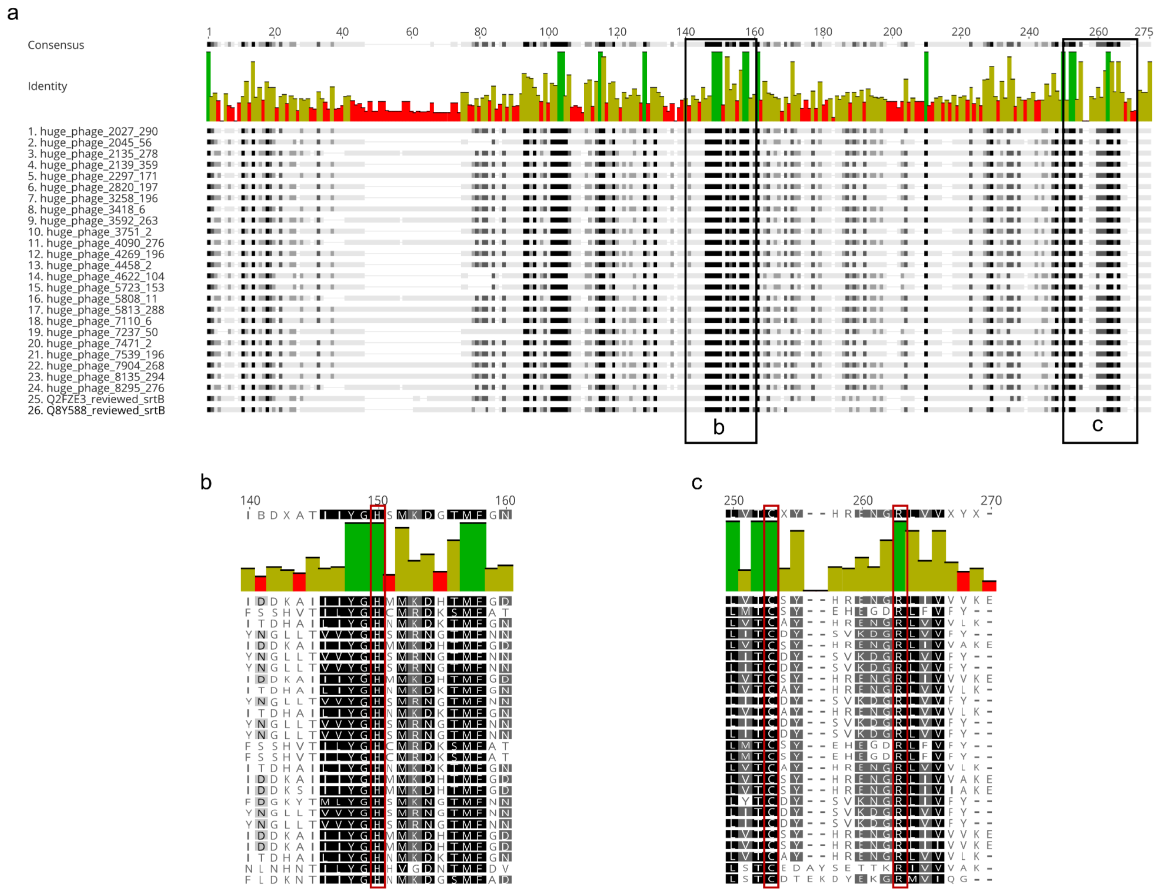


**Supplementary Figure S2 | Conservation of key residues in huge phage-encoded sortase B proteins.** (a) Sequence alignment of 24 huge phage-encoded sortase B (SrtB) proteins and two reviewed bacterial SrtB reference proteins (UniProtKB IDs: Q2FZE3 and Q8Y588). Black boxes indicate the regions displayed in b and c. The sequences were aligned using MUSCLE and visualized in Geneious Prime. The consensus sequence at the top indicates the most dominant amino acid at each aligned position. The truncated alignments in (b) and (c) show regions surrounding the conserved (b) His residue and (c) Cys and Arg residues. Red boxes highlight these three conserved residues.

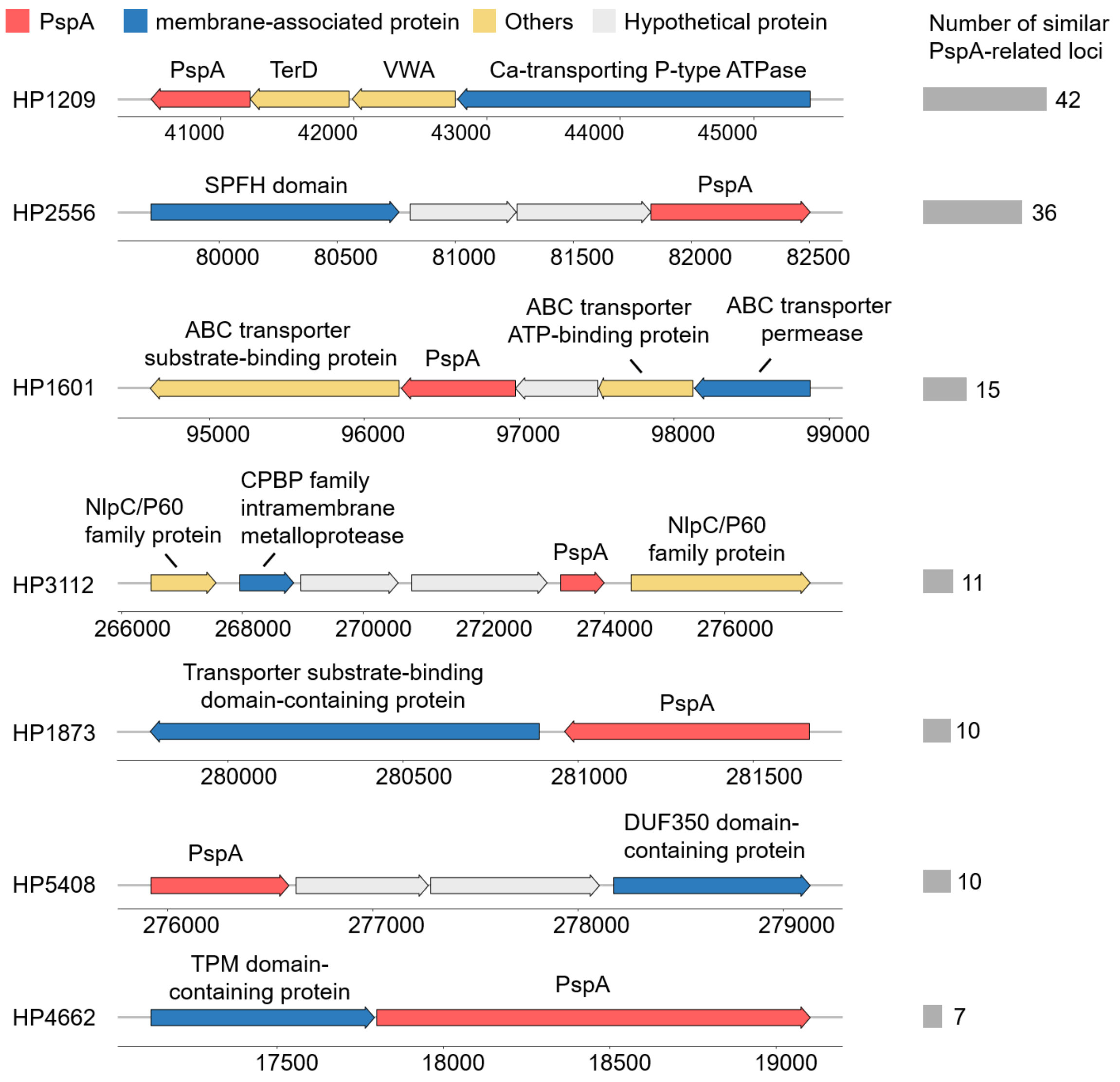


**Supplementary Figure S3 | The genomic neighborhoods of *pspA* in representative huge phage genomes.** Representative genomic neighborhoods containing genes encoding membrane-associated proteins near *pspA* are shown on the left. Arrows indicate genes and their orientations, with *pspA* in red, other membrane-associated protein genes in blue, other annotated genes in yellow, and hypothetical protein genes in grey. All the annotations (if available) are shown above the corresponding genes. Bars on the right indicate the number of *pspA* loci sharing the corresponding encoding pattern, defined by the presence of nearby genes encoding the same membrane-associated protein type.

a

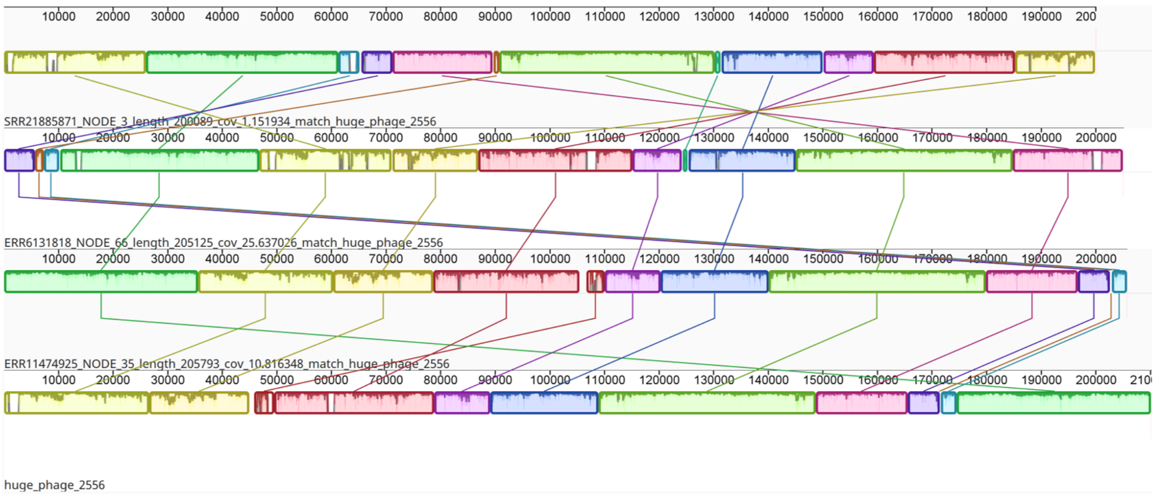


b

| Query genome | Reference genome | ANI (%) | Matched fragments | Total fragments | AF (%) |
|---|---|---|---|---|---|
| ERR6131818_NODE_66_length_205125_cov_25.637026_match_huge_phage_2556 | huge_phage_2556 | 97.5 | 65 | 68 | 95.6 |
| ERR11474925_NODE_35_length_205793_cov_10.816348_match_huge_phage_2556 | huge_phage_2556 | 99.8 | 66 | 68 | 97.1 |
| SRR21885871_NODE_3_length_200089_cov_1.151934_match_huge_phage_2556 | huge_phage_2556 | 96.9 | 64 | 66 | 97.0 |

**Supplementary Figure S4 | The genome-wide comparison of HP2556 and three related genomes.** Three HP2556-related genomes were assembled from SRR21885871(de Nies et al. 2023), ERR6131818 (Hibberd et al. 2024), and ERR11474925 (Cunningham-Oakes et al. 2025), respectively, and used for metatranscriptomic mapping (see Figure 5d in the main text). All three metagenomic samples were collected from the human gut and paired with metatranscriptomic samples (i.e., SRR21885900, ERR8517442, and ERR11492508, respectively). (a) Whole-genome comparison using Mauve within Geneious Prime (Kearse et al. 2012). The homologous genomic regions identified by pairwise sequence alignment were visualized in colored blocks. (b) Whole-genome comparison using FastANI. ANI: average nucleotide identity; AF: Alignment fraction.

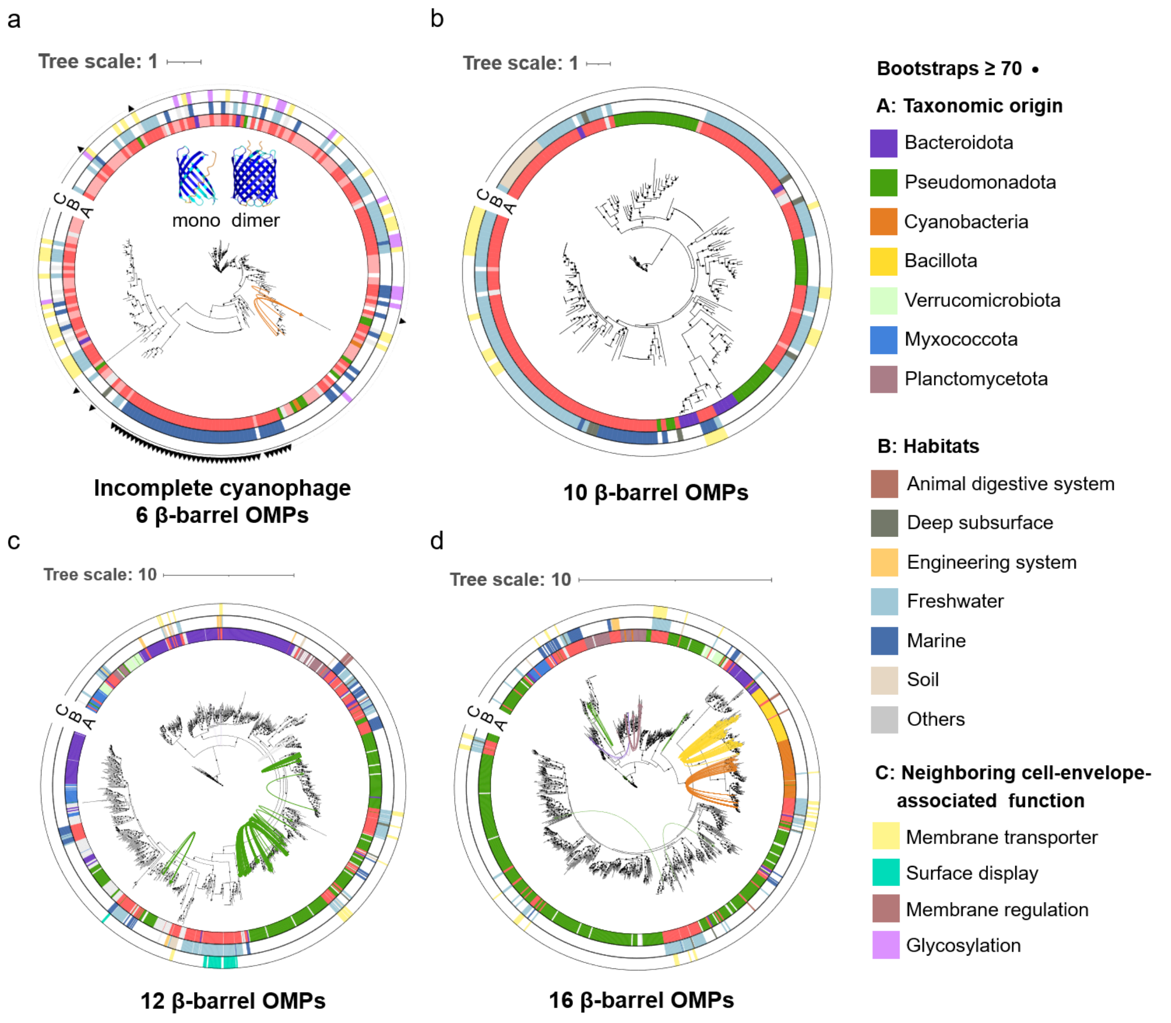


**Supplementary Figure S5 | Phylogenetic relationships and genomic contexts of huge phage-encoded outer membrane proteins.** (a–d) Phylogenetic trees of huge phage-encoded OMPs and bacterial homologs containing (a) 6, (b) 10, (c) 12, and (d) 16 β-strands. Outer color strips indicate taxonomic origin (A), habitat (B), and neighboring gene function (C), respectively. Curves connect huge phage OMPs to homologs in their predicted host genomes. Triangles in (a) mark cyanophage OMPs whose genes lie between genes encoding the major capsid and tail tube proteins. The inset in (a) shows monomeric and dimeric structural models of the six-stranded cyanophage OMP family.

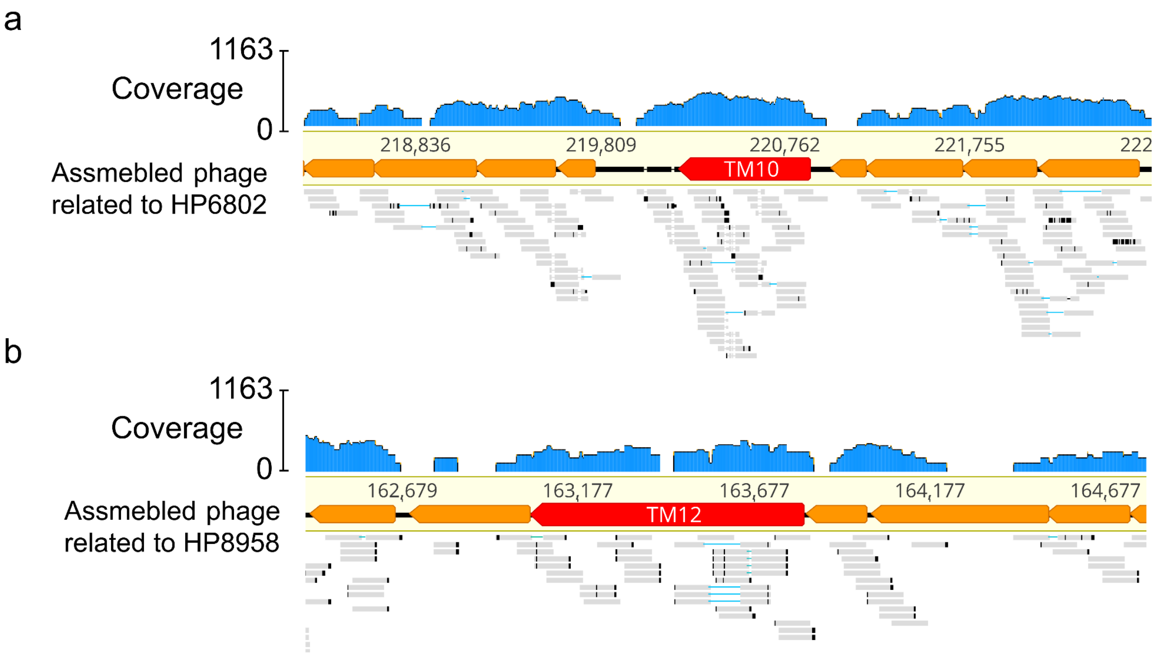


**Supplementary Figure S6 | RNA read mapping profiles showing the transcriptional activity of 10- and 12-stranded outer membrane proteins encoded by huge phages.** (a) Representative genomic region containing a gene encoding a 10-stranded outer membrane protein (TM10). The genome was assembled from a freshwater metagenome (NCBI SRA ID: SRR8437931) and is related to the HP6802 genome cluster. Reads from the corresponding paired metatranscriptome SRR7963492 were mapped to the assembled genome. (b) Representative genomic region containing a gene encoding a 12-stranded outer membrane protein (TM12). The genome was assembled from a marine metagenome (NCBI SRA ID: ERR3589587 (Ngugi et al. 2023)) and is related to the HP8958 genome cluster. Reads from the corresponding paired metatranscriptome ERR3586697 were mapped to the assembled genome. The metagenomic samples were assembled using metaSPAdes (see Methods in the main text for details).
